# The ER-resident Ras Inhibitor 1 (Eri1) of *Candida albicans* inhibits hyphal morphogenesis via the Ras-independent cAMP-PKA pathway

**DOI:** 10.1101/2024.02.28.582498

**Authors:** Subhash Chandra Sethi, Monika Bharati, Usha Yadav, Yatin Kumar, Sneha Sudha Komath

**Author notes:** These authors have contributed equally to the work (Subhash Chandra Sethi, Monika Bharati).

## Abstract

Ras signaling and glycosylphosphatidylinositol (GPI) biosynthesis are mutually inhibitory in *S. cerevisiae*. The inhibition is mediated via an interaction of yeast Ras2 with the Eri1 subunit of its GPI-*N*-acetylglucosaminyl transferase (GPI-GnT), the enzyme catalyzing the very first GPI biosynthetic step. In contrast, Ras signaling and GPI biosynthesis in *C. albicans* are mutually activated and together control the virulence traits of the human fungal pathogen. What might be the role of Eri1 in this pathogen? The present manuscript addresses this question while simultaneously characterizing the cellular role of CaEri1. It is either non-essential or required at very low levels for cell viability in *C. albicans*. Severe depletion of CaEri1 results in reduced GPI biosynthesis and cell wall defects. It also produces hyperfilamentation phenotypes in Spider medium as well as in bicarbonate medium containing 5% CO_2_, suggesting that both the Ras-dependent and Ras-independent cAMP-PKA pathways for hyphal morphogenesis are activated in these cells. Pull-down and acceptor-photobleaching FRET experiments suggest that CaEri1 does not directly interact with CaRas1, but does so through CaGpi2, another GPI-GnT subunit. CaGpi2 is also downstream of CaEri1 in cross-talk with CaRas1 and control of hyphal growth in Spider medium. However, CaEri1 is downstream of all GPI-GnT subunits in inhibiting Ras-independent filamentation. *CaERI1* also participates in the inter-subunit transcriptional cross-talk within the GPI-GnT, a feature unique to *C. albicans*. Virulence studies using *G. mellonella* larvae show that a heterozygous strain of *CaERI1* is better cleared by the host and is attenuated in virulence.

## Introduction

Eukaryotic cell membranes possess a unique class of proteins anchored to the extracellular leaflet *via* a covalently linked glycolipid, the glycosylphosphatidylinositol (GPI) anchor. In organisms that have a cell wall, GPI-APs are also covalently linked to it (1). Universally, the conserved core of the GPI anchor has a phosphatidylinositol (PI) linked to a glucosamine (GlcN), followed by three successive mannose (Man) residues, and an ethanolamine phosphate (EtNP) on the third Man to which the protein is attached *via* an amide bond, to produce PI-GlcN4-1αMan6-1αMan2-1αMan6-EtnP-Protein, in the lumen of the endoplasmic reticulum (ER). A fourth Man α1-2 is compulsorily added to the third Man in fungi (1). Further modifications of this conserved core can occur in a cell-type and species-dependent manner. The types of proteins that post-translationally receive the GPI anchor depend on the species and cell-types in which they are expressed. In *C. albicans*, many of the proteins that receive a GPI anchor are involved in host recognition and virulence (2).

GPI-*N*-acetylglucosaminyltransferase (GPI-GnT) catalyzes the first step of GPI biosynthesis, involving transfer of *N*-acetylglucosamine (GlcNAc) to PI. *S. cerevisiae* GPI-GnT comprises six subunits: Gpi1, Gpi2, Gpi3, Gpi15, Gpi19 and Eri1. Gpi3 (or Spt14) is the likely catalytic subunit, while the others are proposed to be accessory subunits. Yil102c-A, a distant homolog of mammalian DPM2, was recently identified as a seventh subunit, which interacts with Gpi3 on the one hand and with the DPM1 subunit of the dolichol-phospho-mannose (Dol-P-Man) synthase on the other, thus linking GPI biosynthesis with Dol-P-Man synthesis in a manner similar to that seen in mammals (3, 4).

GPI biosynthesis in *S. cerevisiae* also interacts with the Ras signaling pathway *via* the same enzyme complex. The GPI-GnT inhibits, and is in turn inhibited by, ScRas2 (5). The **<u>E</u>**ndoplasmic reticulum-associated **<u>R</u>**as **<u>I</u>**nhibitor **<u>1</u>** (Eri1) subunit of the GPI-GnT is so named because it inhibits Ras signaling by physically binding to ScRas2 in an effector loop dependent manner; its loss results in hyperactive Ras signalling (6). The ScGpi2 subunit co-precipitates with ScEri1 and with ScRas2 (5). Additionally, *Scgpi1Δ*, *Scgpi2Δ* and *Scgpi19Δ* mutants are all hyperfilamentous, and both *Scgpi3Δ* and *Scgpi15Δ* mutants are hypersensitive to heat, suggesting that hyperactive Ras signaling is probably associated with the GPI-GnT as a whole (5–7).

At least three features distinguish the *C. albicans* GPI-GnT from that of *S. cerevisiae*. Firstly, not all the mutants of the GPI-GnT are hyperfilamentous. Heterozygous strains of *CaGPI1, CaGPI2* and *CaGPI3*, are hypofilamentous while those of *CaGPI19* and *CaERI1* are hyperfilamentous (8). This feature is a result of the unique inter-subunit transcriptional regulations within the *C. albicans* GPI-GnT, which do not have a parallel in *S. cerevisiae* (8, 9). Hyphal growth of these mutant strains, in media that induce Ras-cAMP-PKA signaling, correlates with the expression of CaGpi2 (8). Overexpression of CaGpi2 in the wild type strain or in any of the hypofilamentous GPI-GnT mutants activates the Ras/cAMP/PKA signaling pathway and promotes filamentation. Disrupting one allele of *CaGPI2* in heterozygous strains of *CaGPI19* or *CaERI1* brings down CaGpi2 levels and abrogates their hyperfilamentous phenotype observed in glucose or Spider media. Further, *C. albicans* GPI-GnT is activated, rather than inhibited, by CaRas1, a homolog of ScRas2 (8). This involves a physical interaction of CaRas1 with the CaGpi2 subunit at the endoplasmic reticulum (ER) and is favored by the GTP-bound active form of CaRas1. ScRas2 cannot functionally replace CaRas1 at this step, although it complements the function of CaRas1 in the cAMP/PKA signaling cascade at the plasma membrane (PM) (10).

If the CaEri1 subunit does not directly control Ras/cAMP/PKA signaling, then what might be its role in *C. albicans*? The present manuscript seeks to specifically address this question. We demonstrate that *CaERI1* is haplosufficient for growth and is required for GPI-GnT activity. Like in *S. cerevisiae* (6), *CaERI1* is not essential for viability and its depletion induces higher chitin production and thicker cell walls. While Ras signaling is indeed upregulated in CaEri1 deficient strains, this pathway is primarily controlled by CaGpi2. The CaEri1 subunit, on the other hand, controls hyphal morphogenesis primarily *via* a Ras-independent cAMP/PKA pathway. Finally, a heterozygous strain of *CaERI1* is less likely to kill *G. mellonella* larvae as compared to the wild type strain, thus implicating it in the control of virulence traits in *C. albicans*.

## Results

All the plasmids used in this study, as well as the strains used, their genotypes, and the manner in which they are referred to in the text are given in Table-I.

**Table-I:** List of *C. albicans* strains used in this study.

| **A. List of *C. albicans* strains** |  |  |  |  |
| --- | --- | --- | --- | --- |
| S. No. | Strain referred to in text as | Strain Nomenclature | Strain Genotype | Reference |
| 1. | BWP17 | BWP17 | *ura3::imm434/ura3::imm434*<br>*iro1/iro1::imm434*<br>*his1::hisG/his1::hisG arg4/arg4* | (14) |
| 2. | BWP17-URA3 | BWP17-URA3 | BWP17-*rps1Δ::P<sub>ACT1</sub>-GFP::URA3/RPS1* | (26) |
| 3. | *CaERI1Hz* | *Caeri1/CaERI1* | BWP17-*eri1Δ::HIS1/ERI1* | (8) |
| 4. | *CaERI1Hz-URA3* | *Caeri1/CaERI1/P<sub>ACT1</sub>-URA3* | BWP17-*eri1Δ/ERI1::rps1::pACT1-GFP::URA3/RPS1* | (8) |
| 5. | *CaERI1 reintegrand* | *Caeri1/CaERI1/P<sub>ACT1</sub>-CaERI1* | BWP17-*eri1Δ::HIS1/ERI1::RPS1/rps1::pACT1-ERI1::URA3* | (8) |
| 6. | *Caeri1 null* | *Caeri1/ P<sub>MET3</sub>-CaERI1* | BWP17-*eri1Δ::HIS1/P<sub>MET3</sub>-ERI1::URA3* | This study |
| 7. | *Caras1 null* | *Caras1/Caras1* | BWP17-*ras1Δ::HIS1/ras1Δ::ARG4* | (31) |
| 8. | *Caeri1 null/CaRAS1Hz* | *Caras1/CaRAS1/Caeri1/CaERI1* | BWP17-*eri1Δ/ P<sub>MET3</sub>-ERI1::URA3::ras1Δ/RAS1::ARG4* | This study |
| 9. | *Caras1 null/CaERI1Hz* | *Caras1 null/CaERI1Hz* | BWP17-*ras1Δ::HIS1/ras1Δ::ARG4::eri1Δ::HIS1/ERI1* | This study |
| 10. | *CaGPI1Hz* | *Cagpi1/CaGPI1* | BWP17-*gpi1Δ::ARG4/GPI1* | (8) |
| 11. | *CaGPI2Hz* | *Cagpi2/CaGPI2* | BWP17- *gpi2Δ::ARG4/GPI2* | (26) |
| 12. | *CaGPI3Hz* | *Cagpi3/CaGPI3* | BWP17-*gpi3Δ::HIS1/GPI3* | (8) |
| 13. | *CaGPI15Hz* | *Cagpi15/CaGPI15* | BWP17-*gpi15Δ::HIS1/GPI15* | (8) |
| 14. | *CaGPI19Hz* | *Cagpi19/CaGPI19* | BWP17-*gpi19Δ::HIS1/GPI19* | (32) |
| 15. | *CaGPI1Hz/CaERI1Hz* | *Cagpi1/CaGPI1/Caeri1/CaERI1* | BWP17-*gpi1Δ::ARG4 /GPI1::eri1Δ::HIS1/ERI1* | This study |
| 16. | *CaGPI2Hz/CaERI1Hz* | *Cagpi2/CaGPI2/Caeri1CaERI1* | BWP17- *gpi2Δ::ARG4/GPI2 ::eri1Δ::HIS1/ERI1* | (8) |
| 17. | *CaGPI3Hz/CaERI1Hz* | *Cagpi3/CaGPI3/Caeri1/CaERI1* | BWP17- *gpi3Δ::HIS1/GPI3 ::eri1Δ::ARG4/ERI1* | This study |
| 18. | *CaGPI15Hz/CaERI1Hz* | *Cagpi15/CaGPI15/Caeri1/CaERI1* | BWP17- *gpi15Δ::HIS1/GPI15 ::eri1Δ::ARG4/ERI1/* | This study |
| 19. | *CaGPI19Hz/CaERI1Hz* | *Cagpi19/CaGPI19/Caeri1/CaERI1* | BWP17-*GPI19/gpi19Δ::HIS1::eri1Δ::ARG4/ERI1* | This study |
| 20. | *CaGPI2-6X-* | *CaGPI2/CaGPI2-6X-HIS* | BWP17-GPI2/GPI2-6X- | (8) |
|  | *HIS* |  | *HIS::URA3* |  |
| 21. | *CaGPI2-6X-HIS/CaERI1-V5* | *CaGPI2/CaGPI2-6X-HIS/CaERI1/CaERI1-V5* | *BWP17-GPI2/GPI2-6X-HIS::URA3::ERI1/ERI1-V5::ARG4* | This study |
| 22. | *CaERG11-6X-HIS* | *CaERG11/CaERG11-6X-HIS* | *BWP17-ERG11/ERG11-6X-HIS::URA3* | (8) |
| 23. | *CaERI1-V5* | *CaERI1/CaERI1-V5* | *BWP17-ERI1/ERI1-V5::ARG4* | This study |
| **B. List of plasmids used in this study** |  |  |  |  |
| **S. No.** | **Plasmids** | **Reference** |  |  |
| 1. | *pACT1-GFP* | Kind gift from Prof. Alistair Brown (33) |  |  |
| 2. | *pACT1-CaERI1* | (8) |  |  |
| 3. | *pRS-ARG4* | Kind gift from Prof. Aaron P Mitchell (14) |  |  |
| 4. | *pMET3-URA3-GFP* | Kind gift from Dr. Cheryl Gale (16) |  |  |
| 5. | *pTZ-HIS1* | (32) |  |  |

### TMHMM and Multiple sequence alignment analysis (MSA) of CaEri1

The sequence of CaEri1 was obtained from the Candida Genome Database (www.candidagenome.org). *CaERI1* is located on Chromosome 2A as a single copy in SC5314 strain of *C. albicans*. The sequences of ScEri1 and hPIG-Y were retrieved from the NCBI website. *CaERI1* codes for a 116 amino acids long protein whereas its orthologs in *S. cerevisiae* and humans consist of 68 and 71 amino acids, respectively (6, 11). Multiple sequence alignment of the three Eri1 orthologs was done using the T-Coffee server and the result generated using BoxSHADE (12) is shown in Figure 1A. They show poor sequence conservation with ∼3% identity and ∼8% similarity across the three organisms, as analysed by Uniprot Align tool (https://www.uniprot.org/align/). When compared with the individual orthologs, CaEri1 shares ∼14% and ∼16% amino acid identity with hPIG-Y and ScEri1, respectively. Like ScEri1 and hPIG-Y (6, 11), CaEri1 has two putative transmembrane domains as predicted by TMHMM (13), but its predicted orientation more closely resembles that of hPIG-Y than of ScEri1 (Figure 1B (i-iii)).

**Figure 1.**
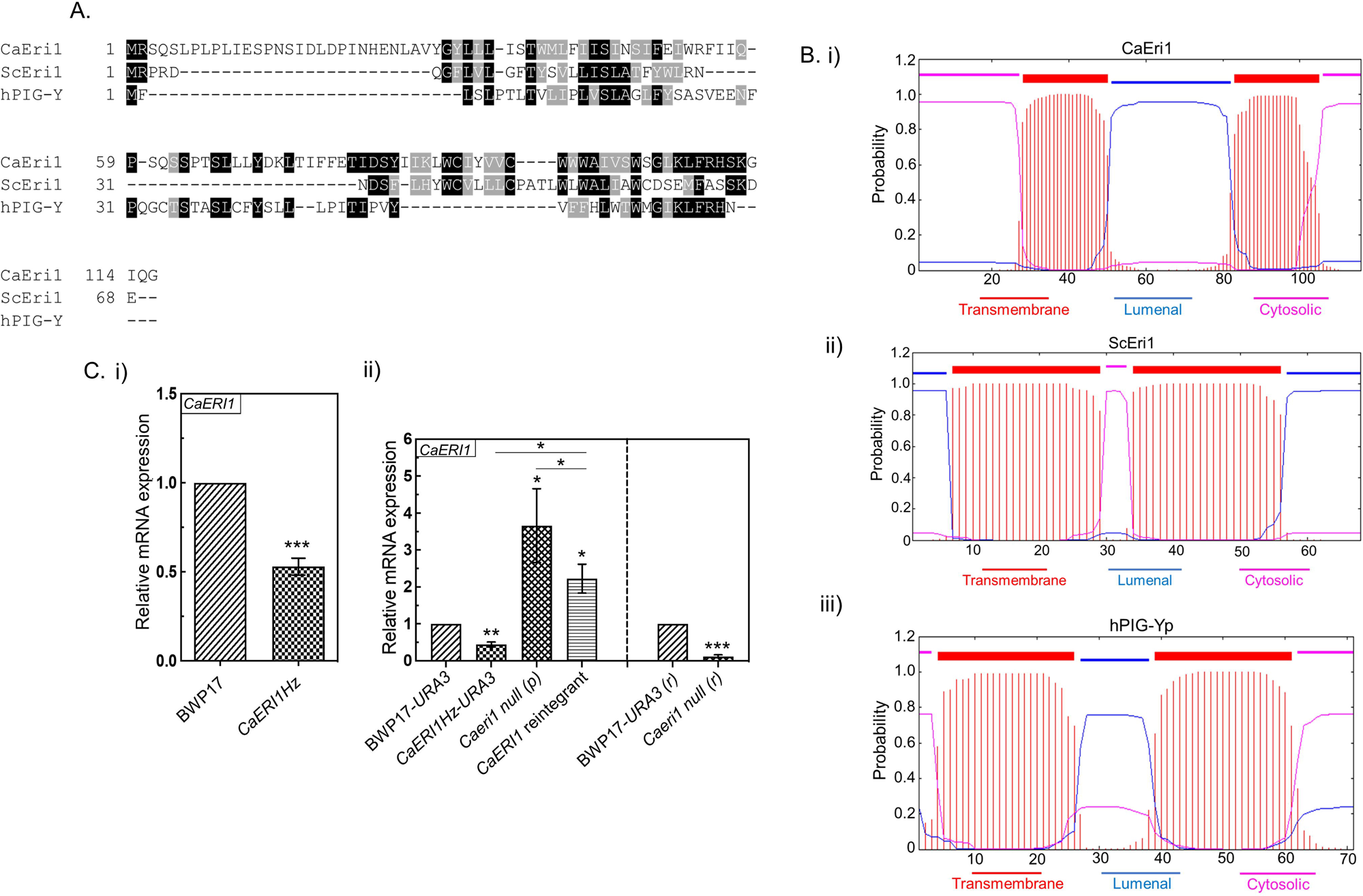
Comparison of CaEri1 with other Eri1 homologs. **A. Multiple sequence alignment of CaEri1 with other Eri1 homologs.** The sequence of CaEri1 was used as Query in BLASTp to identify other homologous proteins. The sequences of CaEri1, ScEri1 and hPIG-Yp were aligned using T-Coffee server and results were visualized using BOXSHADE. The residues in black box represent absolute conservation and the ones with grey show similar types of amino acids. **B. Hydrophobicity profiling using TMHMM.** (i-iii) The amino acid sequences of CaEri1, ScEri1 and hPIG-Y were analysed by TMHMM (http://www.cbs.dtu.dk) for hydrophobicity profiling. **C. Confirmation of *Caeri1* mutant strains by examining the *CaERI1* mRNA levels using real time PCR. (i)** Transcript level analysis of *CaERI1* in *CaERI1Hz* versus BWP17 grown in SD medium containing uridine. **(ii)** Transcript levels of *CaERI1* in *CaERI1Hz-URA3*, *Caeri1 null (p)* and *CaERI1 reintegrant* relative to BWP17-*URA3* grown in SD medium in the absence of Met/ Cys and in *Caeri1 null (r)* versus BWP17-*URA3 (r)* grown in the presence of 1 mM of Met/ Cys. This experiment has been repeated thrice with independent cultures for confirmation. Averages along with standard deviations from three independent sets are plotted.

### Generation of conditional null *Caeri1* strain

Homologous recombination (14) was used to generate a conditional null of *CaERI1* in the BWP17 strain background. As reported previously, *Caeri1/CaERI1* (called *CaERI1Hz* from here on) is a heterozygous strain with one allele of *CaERI1* disrupted with the *HIS1* nutritional marker in a BWP17 strain background (8). A homozygous null strain of *CaERI1* was not obtained despite several attempts. Therefore, a conditional null strain *Caeri1/P_MET3_-CaERI1* (called *Caeri1 null* from here on) was generated with the help of the repressible P*_MET3_* promoter using *URA3* as a selection marker (Figure S1A) (15, 16). Met/Cys represses expression from P*_MET3_* and hence a conditional null strain is obtained under such conditions. When grown in permissive conditions, i.e. in the absence of Met/Cys, this strain will be referred to as *Caeri1 null (p)* and as *Caeri1 null* when grown in the presence of 1 mM Met/Cys (repressive conditions). The reintegrant strain, *Caeri1/CaERI1/*P*_ACT1_-CaERI1* (called *CaERI1* reintegrant from here on) has *CaERI1* placed under the control of the constitutively active P*_ACT1_* promoter at the *RPS1* locus in *CaERI1Hz.* The strains were confirmed by PCR and RT-qPCR (Figure 1C (i-ii); Figure S1B (i-iii)). *URA3* has locus-specific effects on gene expression and phenotypes, but reintegration of a single copy of the gene at the *RPS1* locus can normally compensate for most of these issues (17). Hence, *URA3* integrated at the *RPS1* locus in BWP17 (BWP17-*URA3*) and in *CaERI1Hz* (*CaERI1Hz-URA3*) (Figure S1B (ii)), were used as controls for experiments involving the conditional null and/or reintegrant strains. In all the experiments described below, data for *CaERI1Hz* is compared to that of its wild type control, BWP17. The data for *CaERI1Hz-URA3*, *CaERI1* reintegrant and *Caeri1 null (p)* were compared to BWP17-*URA3* grown in the absence of Met/Cys, while *Caeri1 null (r)* was compared to BWP17-*URA3 (r)* where BWP17*-URA3* is grown in 1 mM Met/Cys.

### Expression of *CaERI1* is not required for growth or viability, but its deficiency alters the cell wall and GPI biosynthesis

*CaERI1Hz* showed no growth defect relative to BWP17 in either liquid SD medium (Figure 2A (i) & (iv)) or on SD-agar (Figure S1C (i)). The *Caeri1 null (p)* and *Caeri1 null (r)* strains too grew normally with no significant growth defects as compared to their respective wild type strains in liquid media (Figure 2A (ii)-(iv)) or on solid media (Figure S1C (ii)-(iii)). No propidium iodide (PI) uptake was observed in any of the strains other than the heat killed positive control (Figure 2B), suggesting that they were all viable. The cell walls of the mutant strains of *CaERI1* exhibited higher staining with calcofluor white (CFW) and Congo red (CR) as compared to the wild type or reintegrant strains (Figures 2C (i)-(iv)); Figure S2).

**Figure 2.**
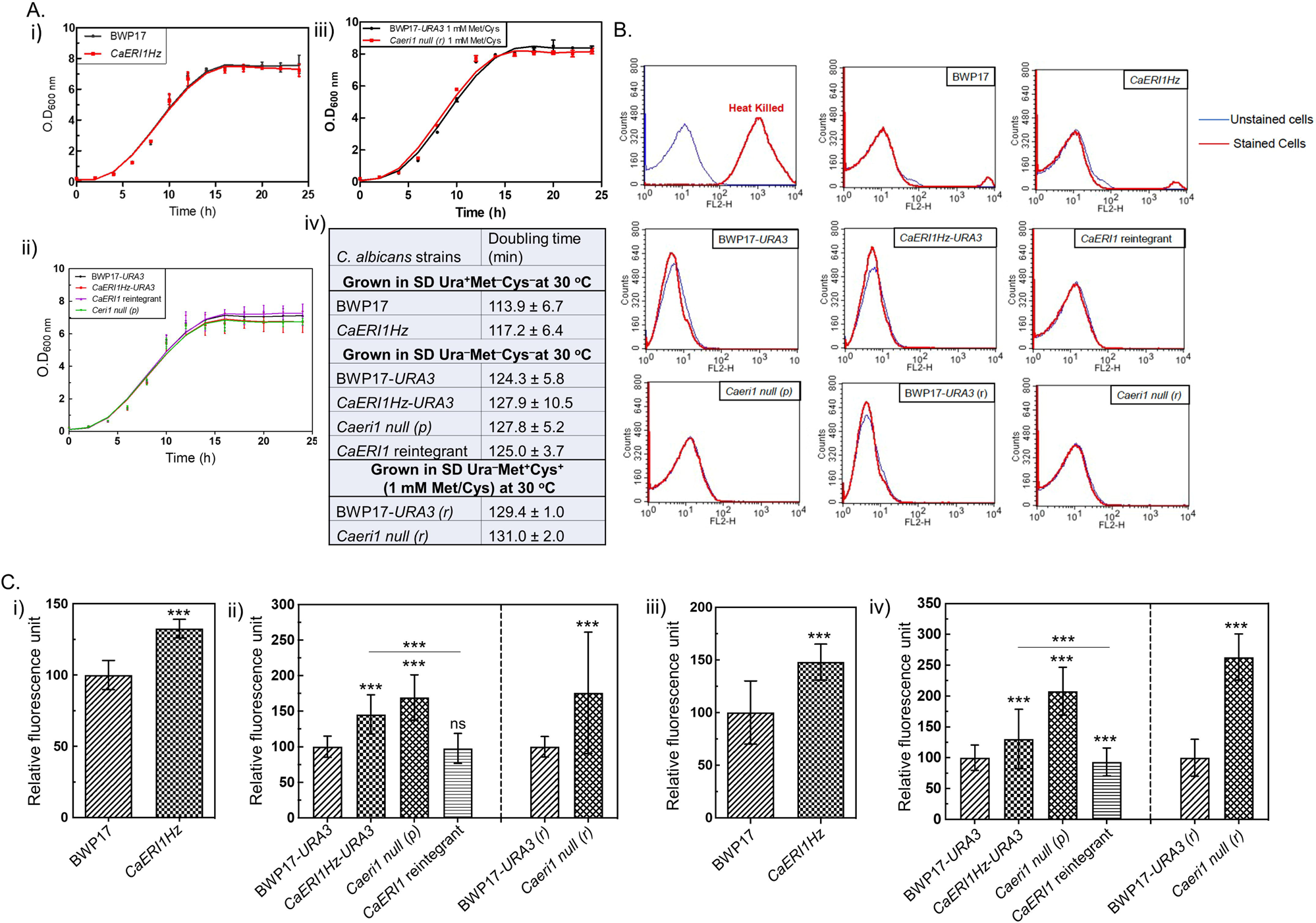
Growth, viability and cell wall defects analyses of *Caeri1* strains. **A. Growth of *CaERI1* strains is unaffected.** The O.D_600 nm_ of each strain grown at 30 °C was monitored at different intervals of time and used to obtain their growth curves. (i) *CaERI1Hz* and BWP17 strains were grown on SD-liquid media containing uridine. (ii) BWP17-*URA3*, *CaERI1Hz-URA3*, *Caeri1 null (p)*, and *CaERI1 reintegrant* strains were grown in SD medium lacking uridine and Met/Cys. (iii) BWP17-*URA3 (r)* and *Caeri1 null (r)* were grown in SD media lacking uridine but containing 1 mM Met/Cys. (iv) The table showing corresponding doubling times of the strains obtained from the growth curves in (i)-(iii). **B. No PI staining is seen in the *CaERI1* deficient strains.** The mid-log phase *Caeri1* mutant strains along with the wild type strains were stained with 5 µg/ml PI. Heat killed BWP17 cells were taken as the positive control. Data corresponding to 1 × 10^5^ cells were acquired for each strain on FL2-H channel. The histograms corresponding to FL2-H (red channel) were overlaid for analysis. **C. *CaERI1* deficient mutants show enhanced staining with CFW and CR**. The strains grown in permissive (p) or repressive (r) conditions were stained with 100 µg/ml CFW (i-ii) or CR (iii-iv) and the fluorescence intensities from 100 cells each were quantified. The data shown are averages with standard deviations of three independent experiments.

GPI biosynthetic activity in the mutant strains was assessed by GPI-GnT assays (Figure 3A) and by measuring the surface levels of Als5, a cell wall localized GPI-AP, by immunostaining (Figure 3B). *CaERI1Hz* exhibited ∼50% reduced GPI-GnT activity as compared to the BWP17 control, while the *Caeri1 null (r)* showed ∼80% reduction in activity relative to BWP17-*URA3 (r)*. In permissive conditions, *Caeri1* is expected to behave like the heterozygous strain, *CaERI1Hz-URA3*, but it showed no significant reduction in GPI-GnT activity vis-à-vis the control strain, BWP17-*URA3* (Figure 3A (i)-(ii)). Nevertheless, this strain showed significant reduction in the levels of Als5 at the cell surface (Figure 3B (iii) & (v)). We consistently saw reduced Als5 expression levels in the heterozygous and conditional null strains (Figure 3B (i)-(v)). GPI-GnT activity was restored in the reintegrant strain relative to *CaERI1Hz-URA3* (Figure 3A (ii)), as were the levels of the GPI-anchored protein Als5 on its cell surface (Figure 3B (iii) & (v)), confirming that the effect was specific to *CaERI1* expression.

**Figure 3.**
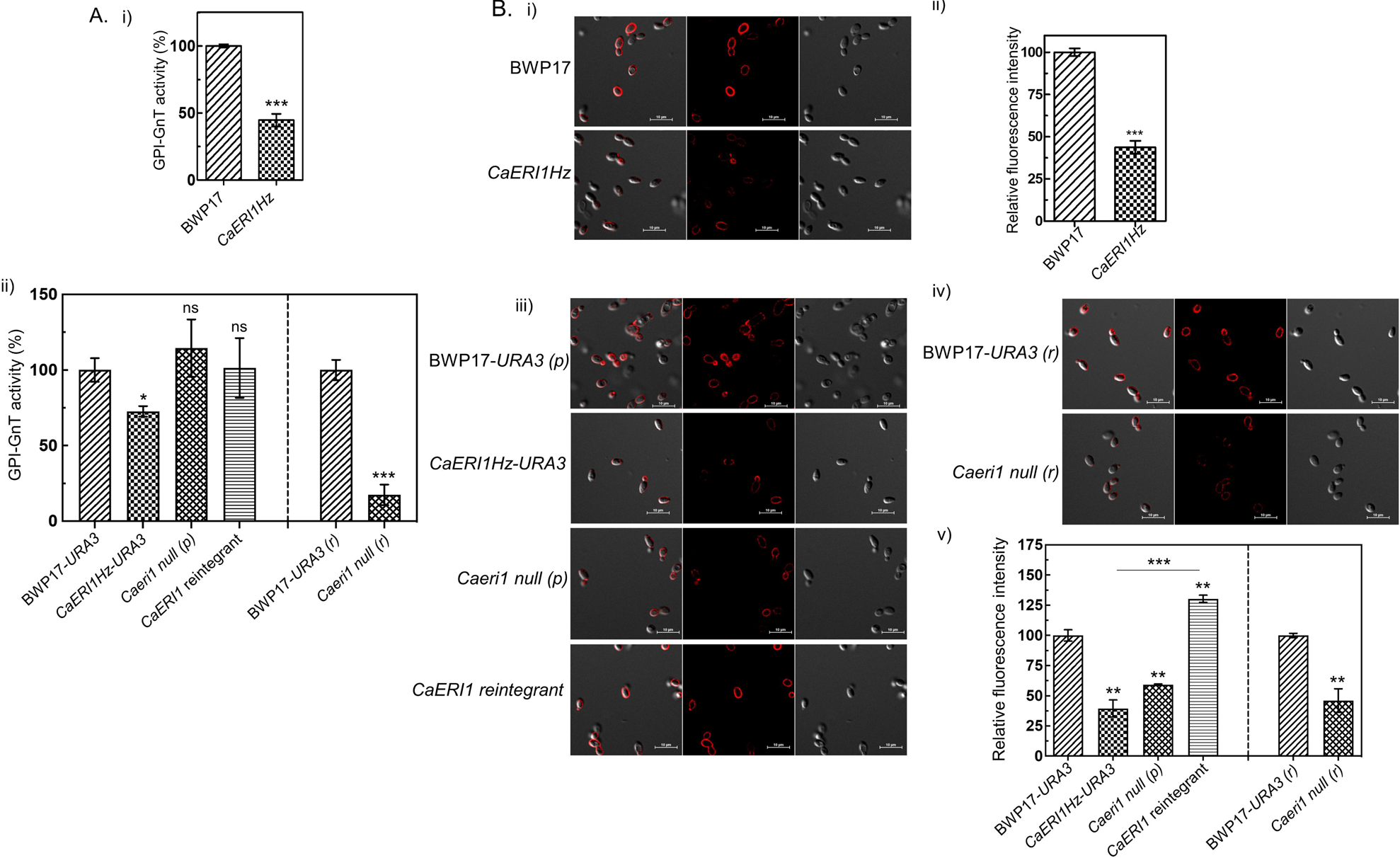
GPI anchor biosynthesis is compromised in *Caeri1* mutant strains. **A. GPI-GnT activity is reduced in the *CaERI1* deficient mutants**. (i-ii) GPI-GnT activity with UDP-[6-^3^H]GlcNAc as the sugar donor was performed using microsomes isolated from strains grown in permissive (p) or repressive (r) conditions. **B. Cell surface expression of GPI-AP is reduced in *CaERI1* deficient cells.** (i, iii & iv) The mid-log phase cells from the mentioned strains were used for immunostaining with anti-Als5 antibody. (ii & v) The bar graphs depict the relative fluorescence intensity (A.U.) of Als5 in the *CaERI1* mutants compared with the respective wild type strains.

### The cAMP-PKA signaling pathway is upregulated in the *CaERI1* mutant strains

The cAMP-PKA pathway is one of the primary dictators of hyphal growth in *C. albicans* (18). This pathway can be induced via Ras-dependent and Ras-independent mechanisms by different hyphae-inducing signals (Figure 4A). For instance, Spider medium induces the former and HCO_3_^−^/ CO_2_ the latter.

**Figure 4.**
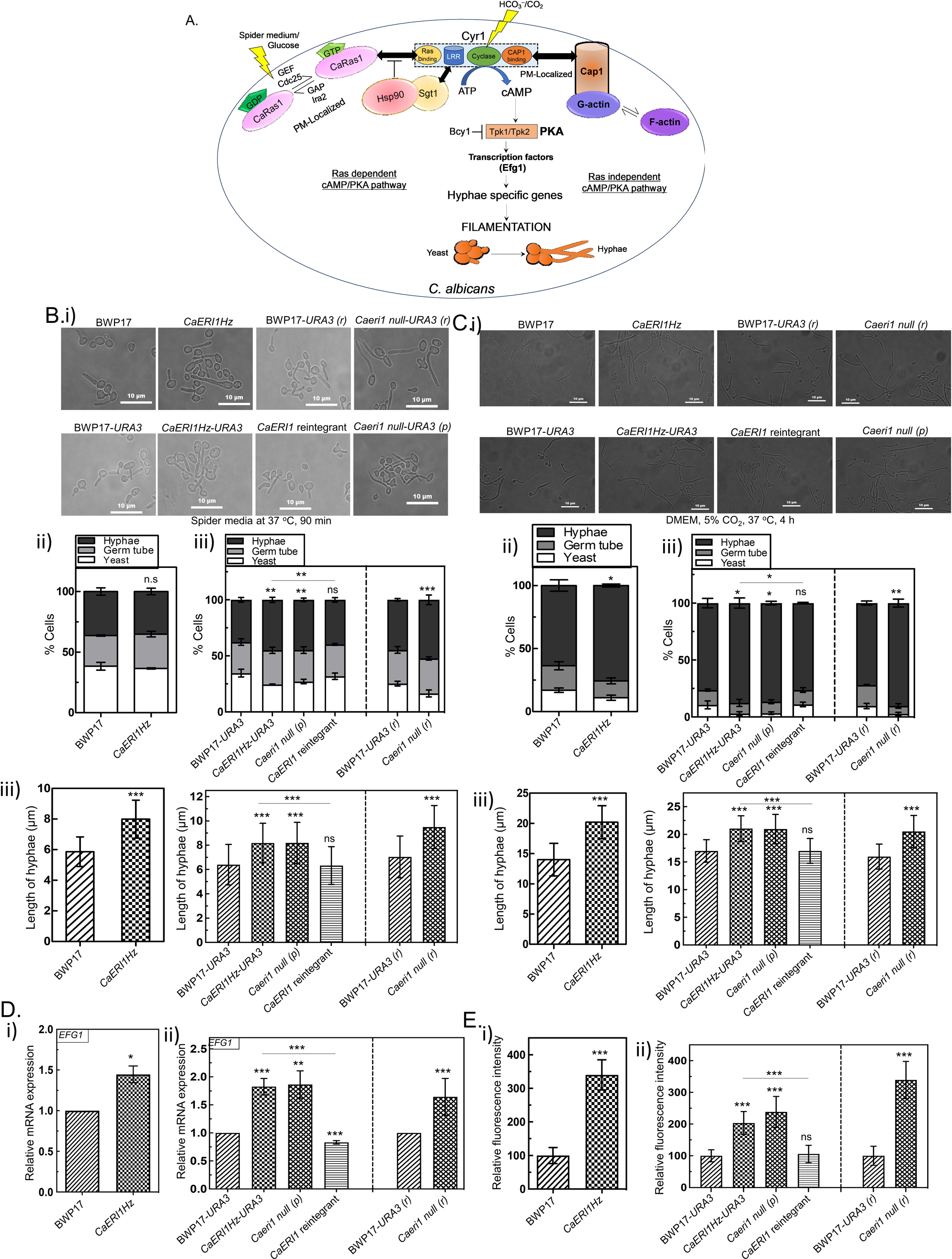
Hyperfilamentation of the *CaERI1* deficient mutants. **A. Schematic diagram showing both Ras-dependent and independent cAMP-PKA pathways.** The cAMP-dependent protein kinase A (PKA) pathway for filamentation can be activated both in a Ras dependent as well as independent manner. Both pathways contribute to filamentation in the *Caeri1* mutants. In response to various stimuli (e.g. Spider medium) inactive CaRas1-GDP is converted to CaRas1-GTP by Cdc25, a guanine nucleotide exchange factor (GEF), which physically interacts with the Ras binding domain of Cyr1, an adenylyl cyclase, to facilitate cAMP/PKA signaling. The heat-shock protein, Hsp90, and its co-chaperone, Sgt1, modulate the interaction. The Ras independent arm of cAMP/PKA pathway is activated either by external stimuli such as HCO ^−^/CO, that are directly sensed by Cyr1, or through increased actin polymerization (F-actin) which is transduced *via* the adenylyl cyclase associated protein 1 (CAP1) binding domain of Cyr1. **B. *CaERI1* deficient cells are hyperfilamentous in Spider medium.** (i) Secondary cultures of the strains were first grown under permissive conditions (Met^−^Cys^−^) and then hyphae were induced in liquid Spider medium at 37 °C for 90 min. Representative images from each mutant strain along with the control are shown. The data shown are averages with standard deviations of (ii) numbers of yeast cells, germ tubes and hyphal cells from each strain counted manually from multiple fields of view and (iii) the lengths of the hyphae in each strain. A total of 80 cells of each strain were used for statistical analysis and the experiment was repeated thrice from independently grown cultures. The statistical significance shown in (ii) are only for hyphal cells. **C. *CaERI1* deficient mutants are hyperfilamentous DMEM + 5% CO2 containing media.** Secondary cultures of the strains were first grown under permissive conditions (Met^−^Cys^−^) and then cultured in DMEM medium at 37 °C in the presence of 5% CO_2_ for 12 h. Representative images for the strains mentioned in the figure are shown. (ii) The bar graphs showing the numbers of yeast cells, germ tubes and hyphal cells from each strain counted manually from multiple fields of view. (iii) Bar graphs depicting the lengths of the hyphae in each strain. A minimum of 80 cells of each strain were used for statistical analysis and the experiment was repeated thrice from independently grown cultures. The statistical significance shown in (ii) are for hyphal cells. **D. Transcript level of *EFG1* increases.** (i-ii) *EFG1* expression at the mRNA level was monitored in *CaERI1* deficient strains versus the wild type controls using RT-PCR. *GAPDH* was used as an internal control for normalization. **E. Actin polymerization is increased in the *CaERI1* deficient mutants.** (i-ii) The F-actin in cells from the strains mentioned in the figure were specifically stained by rhodamine-phalloidin as explained in Experimental Procedures. Fluorescence intensities (arbitrary units) of polymerized actin were quantified and plotted as bar graphs relative to the wild type controls.

*CaERI1Hz* is known to be hyperfilamentous on solid as well as in liquid Spider and YEPD media (8). *CaERI1Hz-URA3* and *Caeri1 null (r)* strains too are hyperfilamentous in liquid Spider medium at 37 °C (Figure 4B (i)). Not only the number of cells producing hyphae but also the average lengths of the hyphae were enhanced in these strains in comparison to BWP17-*URA3* (Figure 4B (ii)-(iii)). Interestingly, the heterozygous as well as conditional null strains of *CaERI1* were also hyperfilamentous in DMEM + 5% CO_2_ at 37 °C (Figure 4C (i)-(iii)). In addition, they exhibited high transcript levels of *EFG1* (Figure 4D), a transcription factor that is activated by cAMP-PKA signaling (19). Phalloidin-staining indicated high levels of F-actin in these strains (Figure 4E; Figure S3). The reintegrant strain showed a reversal of each of these phenotypes. These results suggest that the Ras-dependent and Ras-independent cAMP-PKA signaling pathways are activated in the *CaERI1* deficient mutants.

### CaRas1 is downstream of CaEri1 for filamentation in Spider medium (Ras-dependent cAMP-PKA signaling)

To examine whether CaEri1 was upstream or downstream of CaRas1 or vice versa, two strains *Caras1 null/CaERI1Hz* (where one allele of *CaERI1* is disrupted in a *Caras1/Caras1* homozygous null background) and *Caeri1 null/CaRAS1Hz* (where one allele of *CaRAS1* is disrupted in the *Caeri1 null* strain) were generated along with their appropriate controls (Figure S4 A&B; Table-I) and studied for hyphal morphogenesis in Spider medium as described in Experimental Procedures. As expected, *Caras1 null* shows no hyphal growth (Figure 5A (i)-(ii)) since filamentation in this medium is CaRas1-dependent*. Caras1 null/CaERI1Hz* showed no filamentation either (Figure 5A (i)-(ii)), suggesting that in the absence of any CaRas1, disruption of *CaERI1* does not produce hyperfilamentation. Further, in the *Caeri1 null/CaRAS1Hz* the numbers and length of hyphae are significantly lesser than in the *Caeri1 null* (Figure 5A (i)-(ii)), confirming that CaRas1 acts downstream of CaEri1 in Spider medium.

**Figure 5.**
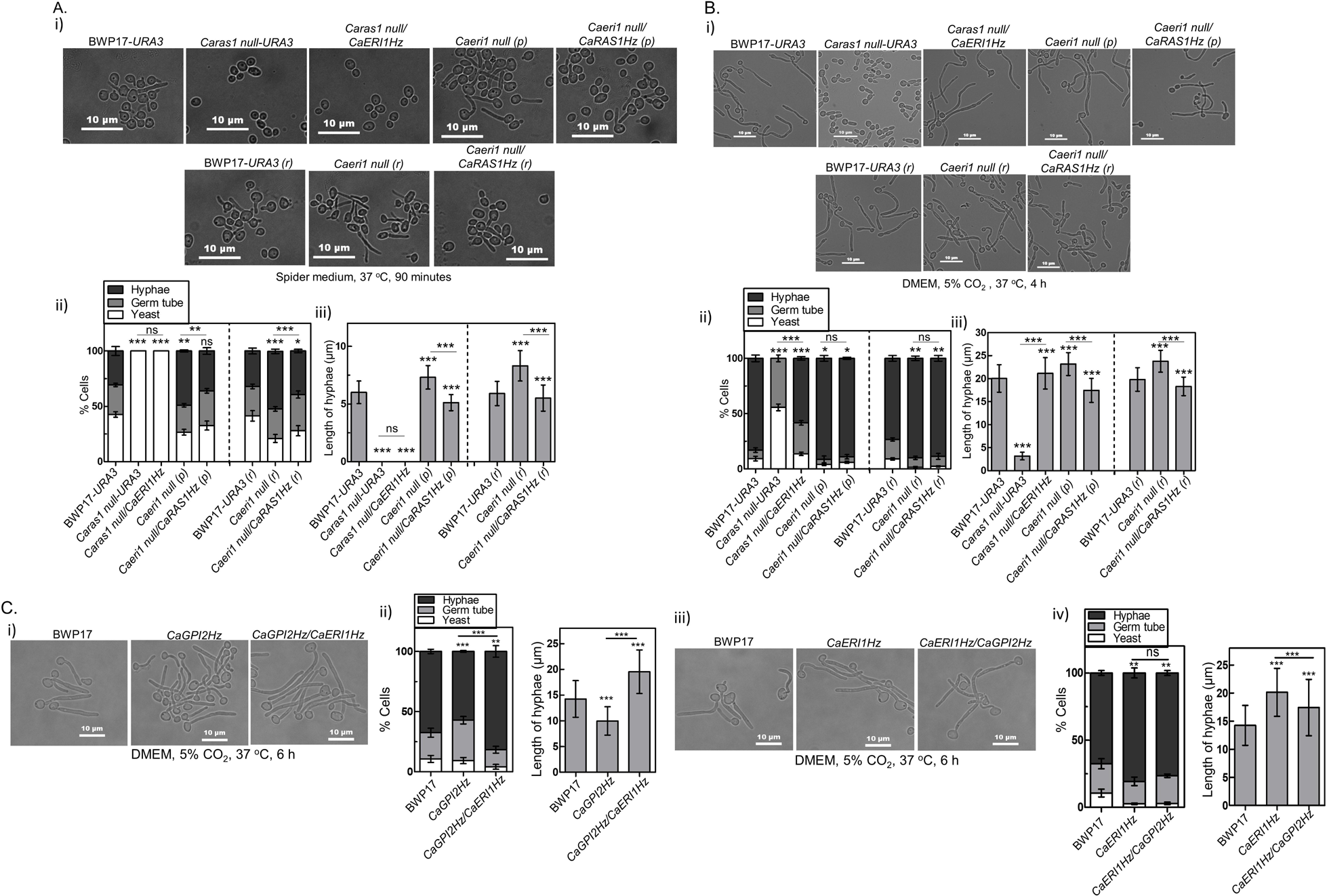
CaEri1 inhibits Ras-independent filamentation. **A. CaRas1 is downstream of CaEri1 for filamentation in Spider medium.** (i) The strains mentioned in the figure were grown in Spider medium for 90 min at 37 °C and representative images are shown. (ii) The average numbers with standard deviations of hyphal, germ tube and yeast cells were plotted as bar chart. The statistical significance shown is for hyphal cells only. (iii) Average hyphal length along with standard deviations are plotted for the same strains. **B. Depletion of CaEri1 induces** filamentation in the presence of HCO ^−^/CO in a CaRas1-independent manner. **(i) The** representative images are shown from the indicated strains when cultured in DMEM medium at 37 °C in the presence of 5% CO_2_. (ii) Average number of cells with standard deviations of different cell types were plotted as bar chart. (iii) Average hyphal length along with standard deviations are plotted for the indicated strains. **C. CaEri1 depletion contributes to increased filamentation in *CaGPI2Hz* strain in response to HCO ^−^/CO**. (i) The mentioned strains were grown in DMEM at 37 °C in the presence of CO_2_ to monitor hyphal growth and the representative images are shown. (ii) Bar graph showing percentage of different cell types in the indicated strains. and hyphal lengths from the same strains were quantified and plotted as bar graph. (iii) Representative images showing filamentation BWP17, *CaGPI2Hz* and *CaGPI2Hz/CaERI1Hz* strains in liquid DMEM medium at 37 °C incubator with 5% CO_2_. (iv) Bar graphs showing percentage of different cell types in the same strains. and hyphal lengths of each strain were also quantified and plotted as bar graph. The statistical significance shown in 720bar graphs depicting the percentage of different cell types in this figure is for hyphal cells only.

### *CaERI1* deficient mutants do not require CaRas1 for filamentation in the presence of HCO ^−^/CO (Ras-independent cAMP-PKA signaling)

To test whether CaRas1 was also downstream of CaEri1 for filamentation in HCO_3_ ^−^/CO_2_, *Caras1 null-URA3* and *Caras1 null/CaERI1Hz* strains (Table-I) were grown in DMEM + 5% CO_2_ as described in Experimental Procedures. Both strains were able to develop hyphae in this medium, although their numbers were lesser than that in the wild type strain (Figure 5B (i)-(ii)). The fact that *Caras1 null-URA3* grows filaments in this medium confirms that hyphal morphogenesis triggered by HCO_3_ ^−^/CO_2_ can occur independently of CaRas1. The basal level of cAMP is expected to be low in this strain which probably explains its reduced filamentation relative to BWP17-*URA3*. *Caras1 null/CaERI1Hz* showed higher numbers of filaments and longer hyphae relative to the *Caras1 null-URA3* strain (Figure 5B (i)-(ii)). Indeed, even in comparison to the wild type strain, the average lengths of hyphae in this strain were marginally higher (Figure 5B (ii)). In contrast, *Caeri1 null/CaRAS1Hz* strain showed no significant decrease in the total numbers but decrease in lengths of hyphae as compared to *Caeri1 null* (Figure 5B (i)-(ii)).

It is already reported that CaGpi2 is downstream of CaEri1 for controlling filamentation in Spider medium (8). To examine whether filamentation in HCO_3_ ^−^/CO_2_ requires CaEri1 to act through CaGpi2, filamentation of two double heterozygous mutant strains, *CaERI1Hz/CaGPI2Hz* (one allele of *CaGPI2* disrupted in a *CaERI1Hz* strain) and *CaGPI2Hz/CaERI1Hz* (one allele of *CaERI1* disrupted in a *CaGPI2Hz* strain), was induced at 37 °C in DMEM + 5% CO_2_. *CaERI1Hz/CaGPI2Hz* showed no significant increase in the numbers of hyphae as compared to BWP17 and a marginal reduction in the average lengths of the hyphae (Figure 5C (i)-(iv)). On the other hand, both these parameters were significantly higher in the *CaGPI2Hz/CaERI1Hz* relative to the *CaGPI2Hz* strain. Thus, disrupting one allele of *CaERI1* activates the HCO_3_ ^−^/CO_2_ mediated Cyr1/cAMP/PKA signaling even in the *CaGPI2Hz* strain.

### Loss of CaEri1 activates HCO_3_ ^−^/CO_2_ mediated filamentation in other GPI-GnT mutants also

Next, we examined whether CaEri1 affects the filamentation of other GPI-GnT mutant strains in media that activates Ras-independent cAMP-PKA signaling. For this, one allele of *CaERI1* was disrupted in heterozygous strains of other GPI-GnT subunits (used in a previous study (8)) giving rise to four additional double heterozygous strains: *CaGPI1Hz/CaERI1Hz*, *CaGPI3Hz/CaERI1Hz*, *CaGPI15Hz/CaERI1Hz* and *CaGPI19Hz/CaERI1Hz* (Table-I). Filamentation in these strains was induced at 37 °C in DMEM + 5% CO_2_. In each case, the percent fraction of hyphal cells and their average lengths were higher than in the parent strains (Figure 6 A-D), suggesting that CaEri1 is downstream of all the GPI-GnT subunits in inhibiting filamentation induced by HCO ^−^/CO.

**Figure 6.**
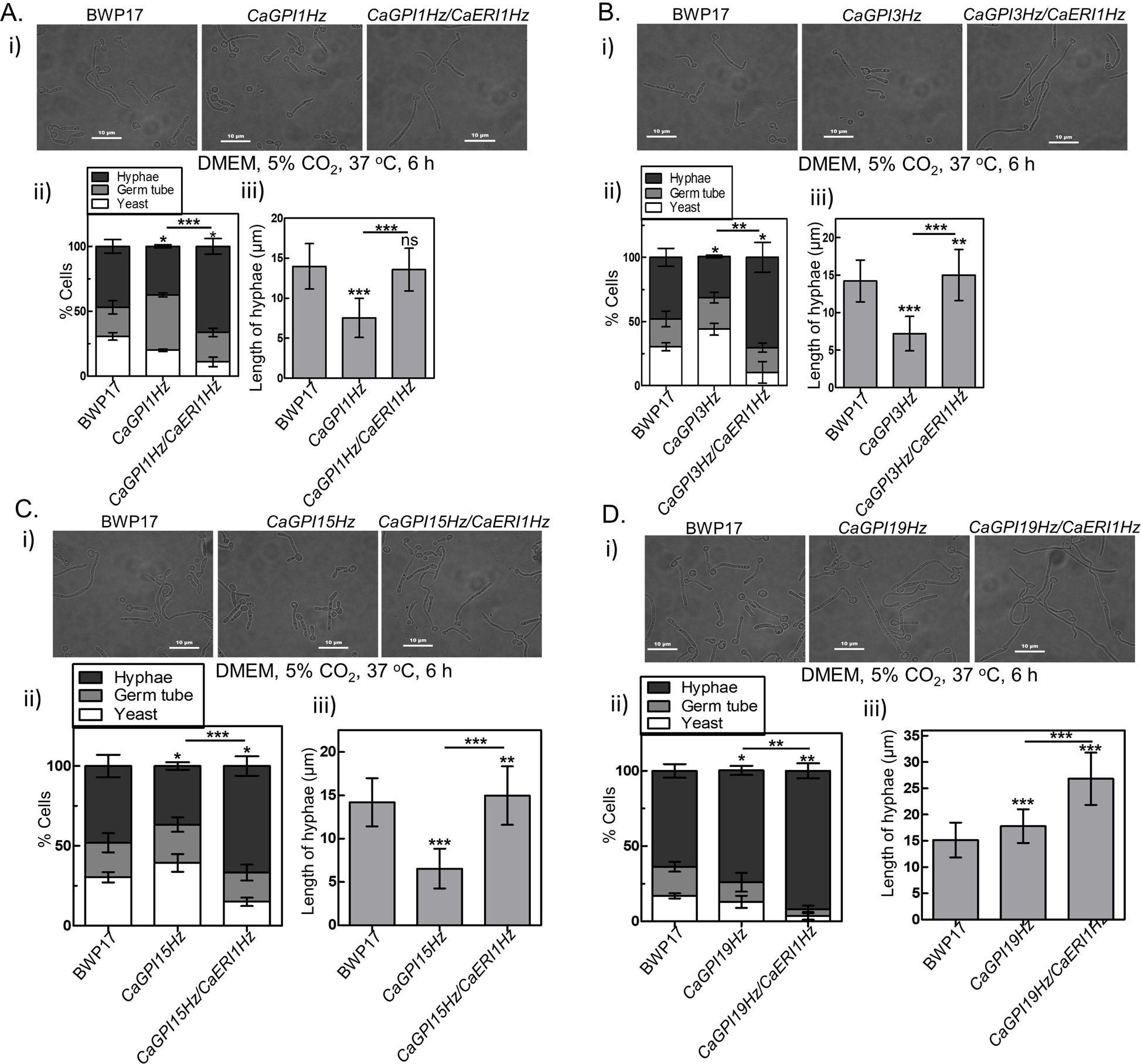
CaEri1 plays a role in controlling filamentation via the Ras-independent pathway in all GPI-GnT mutants. Double heterozygous strains in which one allele of *CaERI1* was disrupted in heterozygous strains of each of the GPI-GnT subunits were studied for their filamentation in HCO ^−^/CO as explained in Experimental Procedures. Shown here are the data for *CaGPI1Hz/CaERI1Hz*(**A**), *CaGPI3Hz/CaERI1Hz* (**B**), *CaGPI15Hz/CaERI1Hz* (**C**), *CaGPI19Hz/CaERI1Hz* (**D**). In each set, (i) Representative images depicting the hyphal growth of the strains when cultured in liquid DMEM medium at 37 °C incubator with 5% CO_2_. (ii) The average percent of yeast, germ tube and hyphal cells from each strain were plotted along with standard deviations. (iii) The average length of hyphae along with standard deviations was plotted as bar graph.

### CaEri1 does not physically interact with CaRas1 but does so with CaGpi2

Since ScEri1 physically interacts with ScRas2 (5), we examined whether a similar interaction might occur in *C. albicans* as well. For this, CaEri1 was C-terminally tagged with V5 in a strain where CaGpi2-6x-His was already expressed (Figure S4C) (for details of the latter strain see (8)). The tagging did not significantly affect the activity of the GPI-GnT complex (Figure 7A). CaEri1-V5 co-localized with the ER marker in the cell (Figure 7B). Both CaEri1 and CaRas1 could be pulled down onto Ni-NTA beads along with CaGpi2-6x-His (Figure 7C). This indicates that the three proteins are part of the same complex.

**Figure 7.**
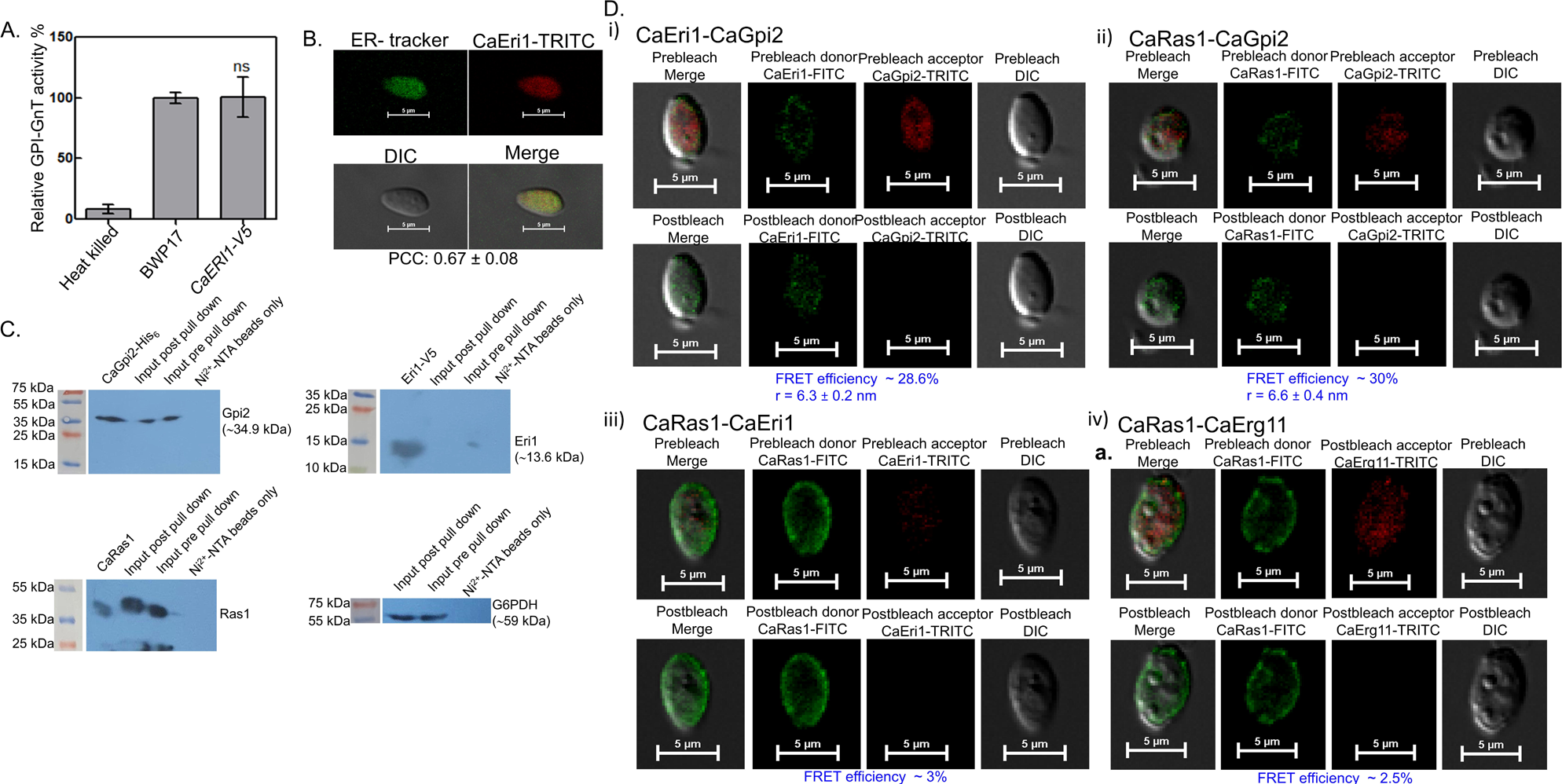
CaEri1 physically interacts with CaGpi2 but not with Ras1. **A. GPI-GnT assay suggests V5-tagging to the C-terminus of Eri1 does not affect the enzymatic activity of GPI-GnT complex.** GPI-GnT activity of the indicated strains was performed by incubating the microsomes with the radiolabelled donor, UDP-[6-^3^H]GlcNAc. **B. CaEri1 localizes to the ER.** Representative immunofluorescence images showing the localization of V5-tagged CaEri1 after immunostained with anti-V5 antibody. ER tracker was used to visualize ER. The Pearson’s correlation coefficient was calculated using the Nikon NIS elements software. **C. Pull-down** assays suggest that CaGpi2, CaRas1 and CaEri1 are part of the same complex. **CaGpi2-His_6_** was pulled down on Ni-NTA beads and run on different SDS-PAGE gels. Western blotting was done using the antibodies mentioned in the figure. **D. AP-FRET suggests that CaEri1 is in close proximity of CaGpi2 but not of CaRas1.** The proteins of interest were probed using primary antibodies followed by appropriate FITC or TRITC labelled secondary antibodies to generate donor-acceptor pairs. Representative AP-FRET images for fluorescence donor-acceptor pairs before and after acceptor photobleaching are shown along with the estimated FRET efficiency and the physical distance between them: (i) CaEri1(FITC)-CaGpi2(TRITC) (ii) CaEri1(FITC)-CaRas1(TRITC) (iii) CaRas1(FITC)-CaGpi2(TRITC), positive control (iv) CaRas1(FITC)-CaErg11(TRITC), negative control.

To determine whether CaEri1 was in close proximity to either CaGpi2 or CaRas1, acceptor photobleaching fluorescence resonance energy transfer (AP-FRET) experiments were performed. As can be seen from Figure 7D (i), upon photobleaching the acceptor, the fluorescence intensity of the donor increased significantly in cells where CaEri1(FITC)/ CaGpi2(TRITC) were present, suggesting that the two proteins are physically close enough to permit energy transfer. AP-FRET was also observed in the positive control, CaRas1(FITC)/ CaGpi2(TRITC) (Figure 7D (ii)). On the other hand, no significant enhancement is seen in the case of CaRas1(FITC)/ CaEri1(TRITC) (Figure 7D (iii)) or in the negative control, CaRas1(FITC)/ CaErg11(TRITC) (Figure 7D (iv)).

### Inter-subunit transcriptional cross-talk within the *C. albicans* GPI-GnT

Inter-subunit transcriptional regulation is a feature unique to *C. albicans* GPI-GnT. From what is known so far, *CaGPI2* and *CaGPI19* are mutually inhibitory and each can independently activate *CaGPI15* expression. CaGpi15 appears to be the master regulator of both *CaGPI2* and *CaGPI19* expressions (20).

To examine how CaEri1 fits in this scheme, we examined the transcript levels of *CaGPI2*, *CaGPI15*, and *CaGPI19* in the *CaERI1Hz* and *CaGPI15Hz* strains (Figure 8A (i)-(ii)). The transcript levels of *CaGPI2* and *CaGPI19* were upregulated in *CaERI1Hz* while that of *CaGPI15* was downregulated. On the other hand, in *CaGPI15Hz*, the expression levels of *CaERI1* were downregulated along with that of *CaGPI2* and *CaGPI19*, suggesting that *CaGPI15* is an activator of all three genes. To confirm this, one allele of *CaGPI15* was disrupted in the *CaERI1Hz* background. The transcript levels of *CaGPI2,* and *CaGPI19* were further reduced in the *CaERI1Hz/CaGPI15Hz* strain (Figure 8A (i)). Disrupting one allele of *CaERI1* in the *CaGPI15Hz* strain background results in upregulation of both *CaGPI2* and *CaGPI19* along with downregulation of *CaGPI15* in the *CaGPI15Hz/CaERI1Hz* strain (Figure 8A (ii)). Thus, *CaERI1* appears to be a repressor of *CaGPI2* and *CaGPI19* and an activator of *CaGPI15.* These results are summarized by the model shown in Figure 8B.

**Figure 8.**
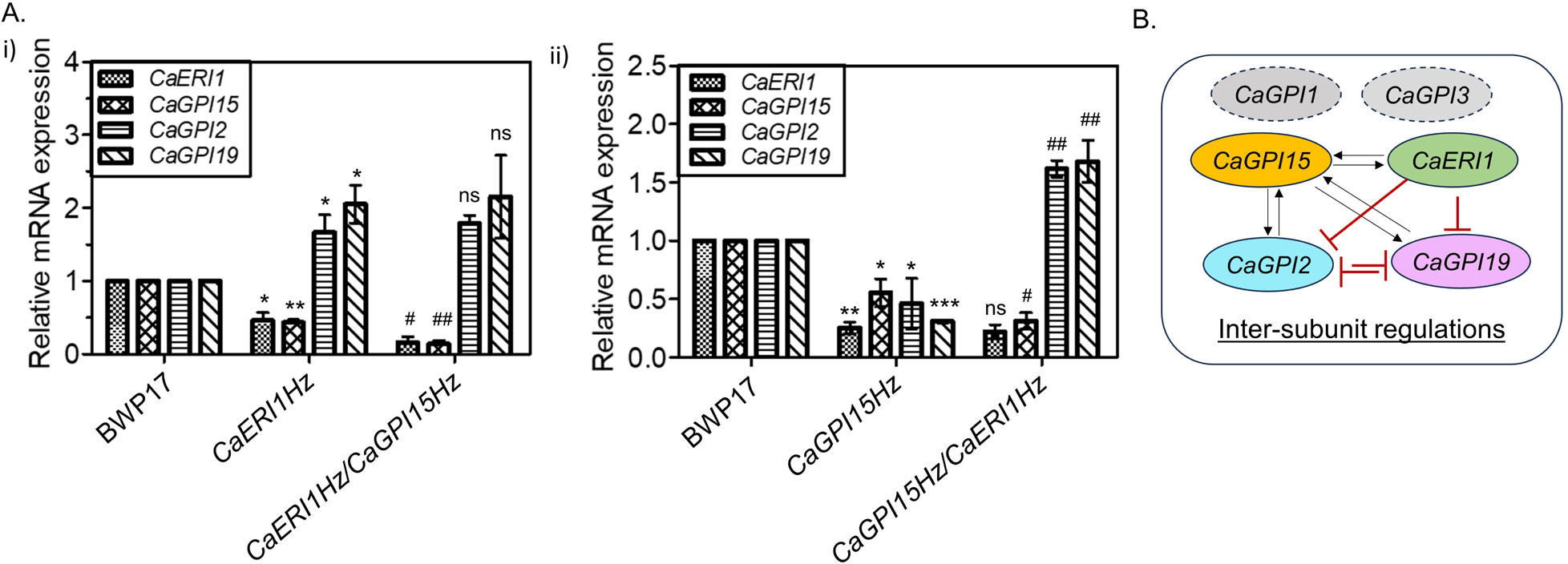
Inter-subunit regulations between members of GPI-GnT complex in *C. albicans*. **A. Transcript levels of GPI-GnT genes in double heterozygous strains.** (i) Transcript levels of *CaERI1*, *CaGPI15*, *CaGPI2* and *CaGPI19* in *CaERI1Hz* and *CaERI1Hz/CaGPI15Hz*. (ii) Transcript levels of *CaERI1*, *CaGPI15*, *CaGPI2* and *CaGPI19* genes in *CaGPI15Hz* and *CaGPI15Hz/CaERI1Hz* strains. **B. The proposed inter-subunit regulations within the GPI-GnT.** *CaERI1* appears to be a transcriptional repressor of *CaGPI2* and *CaGPI19*. *CaGPI15* appears to be a master regulator of all the genes shown. The experiments were performed three times with independent cultures. Note: *P*-values of data for heterozygous strains are shown relative to the wild type strain (represented as *); *P*-values of data for double deletion strains are determined relative to the respective heterozygous parent control (represented as ^#^).

### *CaERI1Hz* cells show reduced virulence in *G. mellonella*

*G. mellonella* larvae are a useful model system for testing the virulence of *C. albicans* (21). Since it would be difficult to ensure gene repression inside the organism for the conditional null strain, the heterozygous and reintegrant strains alone were used in the virulence assays along with the wild type control. The colony forming units (CFU) of *CaERI1Hz-URA3* mutant after 24 h incubation were significantly lesser than that of BWP17-*URA3* (Figure 9A), indicating that the cells of the mutant strain were more susceptible to killing by the host. No CFU were obtained in the PBS injected larvae as well as in the untouched larvae (data not shown). Further, as can be seen from the survival plots (Figure 9B), cells of *CaERI1Hz-URA3* were unable to kill the larvae to the same extent as BWP17-*URA3*. This effect was reversed in the *CaERI1 reintegrant* strain, demonstrating that the effect was gene-specific. No mortality was observed in the control groups.

**Figure 9.**
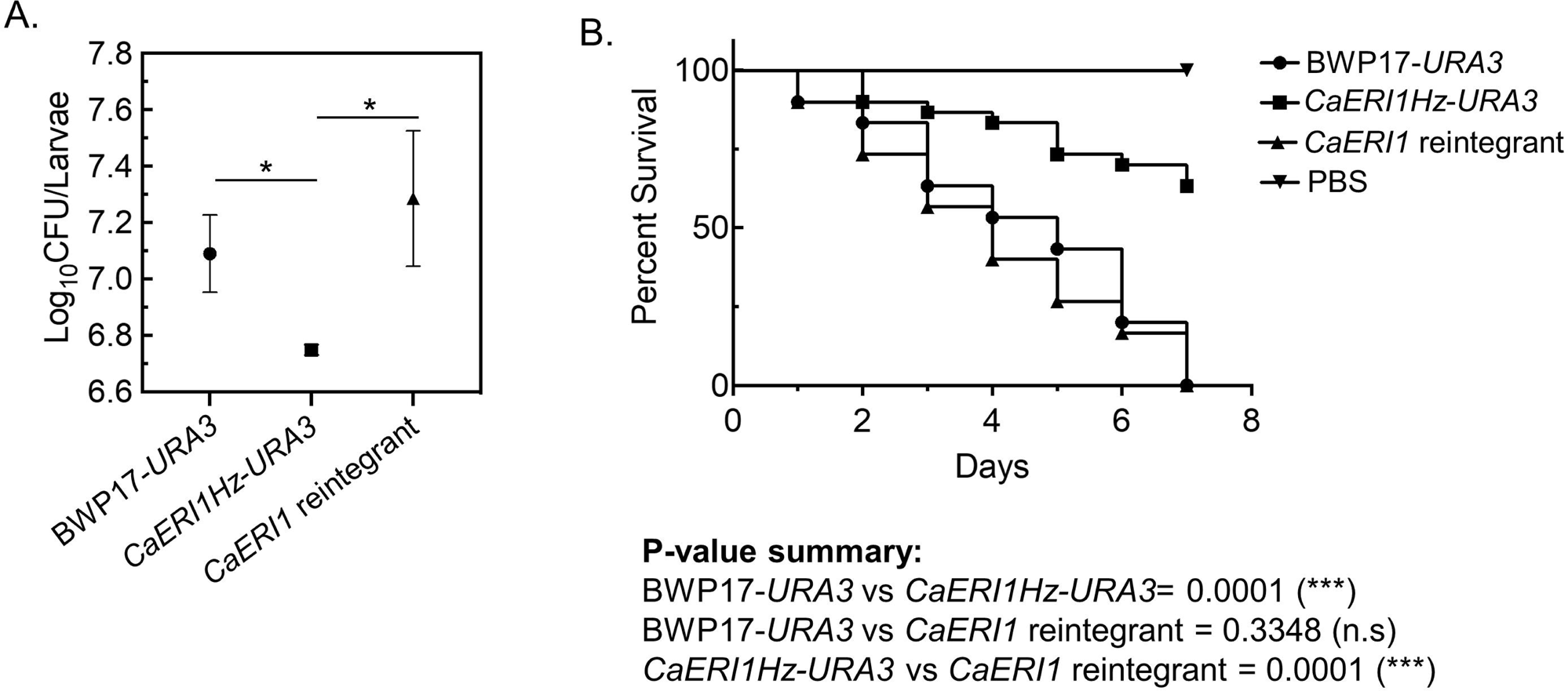
Virulence assay using a *G. mellonella* model. **A.** Fungal burden assay was done by inoculating in each larva 1.0×10^6^ cells of the strains mentioned in the figure. CFU were calculated and log_10_(CFU/larva) was plotted for each. Averages along with standard deviations are shown. **B.** Ten *G. mellonella larvae* were inoculated with 5×10^5^ cells from each strain. Larvae injected with only sterile PBS were taken as controls. Percentage of larvae surviving at the end of each 24 h was plotted.

## Discussion

GPI biosynthesis in all eukaryotes leads to formation of a complex glycolipid anchor with a conserved core consisting of PI-GlcN4-1αMan6-1αMan2-1αMan6-EtnP. The many sequential steps and the enzymes required to build this core structure, is surprisingly more varied than would be expected. In terms of functional conservation of accessory subunits or the regulations of the biosynthetic steps, species-specific differences appear to be even greater and begin right at the very first step of the pathway.

The Eri1/ PIG-Y subunit of the GPI-GnT is perhaps the least conserved of all the subunits of the first enzyme complex. Human PIG-Y and ScEri1 are each required for enhancing endogenous GPI-GnT activity (11). They have roughly 22% sequence identity conserved and two hydrophobic transmembrane segments. But ScEri1 cannot functionally replace hPIG-Y in Daudi cells (derived from human Burkitt lymphoma) as reported in the above manuscript. CaEri1 too has poor homology with respect to either ScEri1 or hPIG-Y, and its predicted membrane orientation resembles that of hPIG-Y rather than of ScEri1. Nevertheless, its endogenous role and the phenotypes of *CaERI1* deficient mutants described here appear to match that of *Sceri1* cells (5, 6). The lack of any significant staining by propidium iodide suggests that *CaERI1* is not required for viability or that low doses of it in the cell will suffice. Repressing its expression in the conditional null *Caeri1* strain significantly reduces GPI-GnT activity and the surface expression of Als5 protein, but does not affect either cell growth or viability. It does, however, attenuate its virulence in a *G. mellonella* infection model.

As explained in the Introduction, the major motivation of this study was to examine whether CaEri1 participates in cross-talk with CaRas1 and whether it plays a role in filamentation of *C. albicans*. Both *Sceri1* mutant cells (6) and *Caeri1* cells are hyperfilamentous. ScEri1 as well as ScGpi2 physically interact with ScRas2 and this has a mutually inhibitory effect on GPI-GnT activity as well as Ras signaling (5). In *C. albicans*, CaGpi2 physically interacts with CaRas1 (the ScRas2 homolog), but the consequence is mutual activation (8). One possible hypothesis would be that CaEri1 too physically interacts with CaRas1 but inhibits it. Pull-down studies indicate that CaRas1 and CaEri1 are in a complex with CaGpi2. However, AP-FRET indicates that CaEri1 and CaRas1 are not positioned close enough for a direct physical interaction. Instead, CaEri1 likely interacts with CaGpi2, which in turn associates with CaRas1.

Multiple experiments in this study demonstrate that the Ras-dependent cAMP/PKA pathway is activated in mutants deficient in *CaERI1*. However, the hyperfilamentous phenotype in Spider medium is primarily controlled by CaGpi2, confirming that the activation of the Ras-dependent cAMP-PKA pathway depends on CaGpi2 rather than CaEri1 (8). However, as also depicted in Figure 4A, *C. albicans* also turns on its cAMP-PKA pathway in a Ras-independent manner, with Cyr1 (the adenylate cyclase), working as a common hub for both arms of the pathway (18). While CaRas1 interacts with the Ras-binding domain of Cyr1 and activates it, CO_2_ can be directly sensed by the cyclase domain of Cyr1 to initiate production of cAMP (22). In a strain completely lacking expression of CaRas1, HCO_3_ ^−^/CO_2_ induced filamentation is higher when one allele of *CaERI1* is disrupted.

Disrupting one allele of *CaGPI2* in a *CaERI1Hz* strain background does not reverse the hyperfilamentous phenotype of the parent strain in CO_2_. On the other hand, disrupting one allele of *CaERI1* in a *CaGPI2Hz* strain switches the parent strain from a hypofilamentous to a hyperfilamentous phenotype under the same conditions. This suggests that CaEri1 controls the Ras-independent arm of the cAMP/PKA pathway, not CaGpi2. It also appears that CaEri1 is downstream of all the other GPI-GnT subunits, and functions as an inhibitor of the HCO_3_ ^−^/CO_2_ induced cAMP-PKA signaling.

For hyphal morphogenesis, as shown in Figure 4A, Cyr1 binds to the CAP1-G-actin complex *via* its CAP1-binding domain and triggers actin polymerization (23). Conversely, polymerization of actin can hyperactivate the cAMP-PKA pathway and produce hyperfilamentation (24). In mutants of *CaERI1* we see increased F-actin levels in yeast cells, indicating that they are already primed for filamentation. Thus, it appears that Eri1 in *C. albicans* has evolved to take on a role different from that in *S. cerevisiae*. Instead of functioning as a Ras-inhibitor, it works as an inhibitor of Ras-independent cAMP-PKA signaling, perhaps directly via Cyr1.

Taken together with our previous results on CaGpi2 (8), it appears that at least two subunits of the *C. albicans* GPI-GnT have evolved to interact with the cAMP-PKA pathway and modulate filamentation in this organism. While the former activates cAMP production via a Ras-dependent pathway, the latter represses it in a Ras-independent manner. They each respond to a different subset of environmental cues typically seen in host tissue environments, enabling GPI biosynthesis in this pathogen to rapidly coordinate with filamentous growth during infection. Inter-subunit transcriptional regulations further aid this process. Overexpression of *CaGPI2* downregulates *CaERI1* expression and *vice versa*, enabling the Ras-dependent and Ras-independent arms to turn on simultaneously, irrespective of which arm of the pathway is first activated.

Thus, it is increasingly evident that the accessory subunits of the GPI-GnT in *C. albicans*, are the crucial hubs through which the GPI biosynthetic pathway cross-talks with and coordinates response to external signals, enabling it to rapidly adapt to the challenges it encounters within the host. Controlling this first step will not only modulate the entire GPI biosynthetic pathway and the expression of GPI anchored proteins at the cell surface but also simultaneously alter the cell wall architecture, sterol biosynthesis and hyphal morphogenesis of this pathogen, thereby attenuating its virulence.

## Experimental Procedures

### Materials

Growth media components were purchased from Hi-Media (India), amino acids, dextrose, galactose, and other routine chemicals from Sisco Research Laboratories (SRL), and glass beads from Unigenetics Instruments Pvt. Ltd. UDP[6-^3^H]GlcNAc was purchased from American Radiolabeled Chemicals Inc. (USA), tunicamycin, syringe filters, HPTLC plates, ECL kits and absolute ethanol were from Merck-Millipore (Germany), DNA gel extraction kit from Qiagen (India), yeast transformation kit from G-Biosciences (USA), trizol reagent and DMEM from Gibco (USA), PCR reagents from Genei (India) and restriction enzymes either from New England Biolabs (USA) or Fermentas (USA). Rhodamine phalloidin was obtained from Invitrogen (Thermo Fisher Scientific, USA). Primers were custom synthesized by GCC Biotech (India), Eurofins Genomics (France) or Sigma-Aldrich, USA. ER tracker BODIPY FL GL was purchased from Thermo Fischer Scientific (USA).

### Strains and growth conditions

The primers used in this study are listed in Table S1. All *C. albicans* strains were generated in BWP17 and grown at 30 C. Ura− strains were grown either in YEPD or in minimal medium containing 60 µg/ml uridine. *Caeri1* conditional null mutants were grown either in permissive (Met− Cys−) or in repressive growth conditions with 1 mM of methionine and cysteine (Met^+^ Cys^+^).

### PCR based amplification of *CaERI1* gene

The forward and reverse primers designed using the sequence retrieved from CGD were used to amplify *CaERI1* from the genomic DNA of BWP17 strain (Table S1). The amplicon (∼ 0.35 kb) was visualized on 1.2% agarose gel.

### Cell viability assays

Propidium iodide (PI) staining was performed as previously described (25). Briefly, primary cultures of the desired strains were grown as above. Equal numbers of cells from secondary cultures (based on OD_600nm_ corresponding to 1.0) were resuspended in PBS, stained with 5 µg/ml propidium iodide for 20 min on a rocker shaker (room temperature, dark), washed thrice with PBS and resuspended in the same buffer. Heat killed wild type cells were taken as the positive control. Flow cytometry was performed on BD FACS Calibur^TM^ (BD Biosciences, California, USA) and 1,00,000 events per sample were recorded. The data were analyzed by BD Cell-Quest Pro software.

### Microsomes preparation from *C. albicans* and GPI-GnT assay

Isolation of microsomes (RER fractions) and GPI-GnT assay were carried out using the standardized protocol from our lab (26). Briefly, the microsomes corresponding to 1.5 mg total protein were incubated in the presence of radiolabelled UDP-[6-^3^H]GlcNAc (1 µCi) and tunicamycin (0.25 mg/ml) for 1 h at 30 °C. Heat killed microsomes were taken as negative controls. The glycolipids were extracted using a solvent mixture having 10:10:3 chloroform/methanol/water followed by drying under N_2_ gas (Grade I). The lipids were then resuspended with water-saturated *n*-butanol and water at 2:1 ratio and the upper butanol layer was collected and dried. Approximately, 10 _μ_l of water-saturated *n*-butanol was used to dissolve the glycolipids and of which, 2 _μ_l was spotted on HPTLC plates and resolved using 65:25:4 chloroform/methanol/water. The radiolabeled glycolipids were detected by Bioscan AR-2000 TLC scanner and quantified using WinScan software.

### Quantification of Als5 levels at the cell surface

Anti-Als5 antibody production was outsourced (27). Primary cultures of the strains were grown on SD medium at 30 °C for 16 h. A 2% inoculum of these was taken to set up secondary cultures and grown for 6 h at 30 °C. Cells from the secondary cultures at O.D_600 nm_ corresponding to 1were taken and incubated with anti-Als5 antibody (1:100) overnight at 4 °C, washed thrice with 1X PBS buffer, and incubated with TRITC-labelled anti-rabbit secondary antibody (1:100) in dark at room temperature for 2 h. Then washed again three times with 1X PBS buffer and resuspended in 50% glycerol. A 10 µl suspension was spotted on glass slides and visualized under a Nikon A1R HD25 confocal microscope. The data were quantified by Nikon NIS element 4000 AR analysis software.

### Microscopy

Where required, cells were spotted on glass slides and visualised under a Nikon Eclipse TiE microscope. The data were quantified by Nikon NIS element 4000 AR analysis software.

### Filamentous growth

Primary cultures of BWP17, *CaERI1Hz, CaGPI2Hz, CaERI1Hz/CaGPI2Hz, CaGPI2Hz/CaERI1Hz, CaGPI3Hz, CaGPI15Hz, CaGPI19Hz, CaGPI3Hz/CaERI1Hz, CaGPI15Hz/CaERI1Hz* and *CaGPI19Hz/CaERI1Hz* were grown overnight at 30 °C in SD media containing uridine; BWP17-*URA3*, *CaERI1Hz-URA3, Caeri1 null (p), CaERI1 reintegrant, Caras1 null-URA3* and *Caeri1 null/CaRAS1Hz (p)*were grown in SD media in the absence of uridine and without Met/Cys (Ura^−^Met^−^Cys^−^); BWP17-*URA3(r)*, *Caeri1 null (r)* and *Caeri1 null/CaRAS1Hz (r)* were grown in SD media containing no uridine but with 1 mM Met/Cys (Ura^−^Met^+^Cys^+^). For hyphal growth in Spider media, 2% inoculum from a primary culture was used for inoculation and hyphae were monitored after 90 min of incubation at 37 °C. For CO_2_ mediated filamentation, 4 ml of Dulbecco’s Modified Eagle Medium (DMEM) containing 40 mM sodium bicarbonate was taken in a 60 mm petri dish, inoculated with 2.5% inoculum from primary culture, and placed at 37 °C in an incubator supplied with 5% CO_2_ for different periods of time. The number of morphological forms (yeast, germ tubes (includes pseudohyphae) and hyphae) were counted manually.

### Cell wall staining with Calcofluor white (CFW) and Congo red

Primary culture of the strains was grown on SD medium at 30 °C for 16 h. From the primary culture, 2% inoculum was taken to set the secondary culture and grown for 6 h at 30 °C. An O.D_600 nm_ corresponding to 0.6 from the secondary cultures was taken to stain the cells either with 100 µg/ml CFW or Congo Red for up to 30 minutes on the rocker shaker under dark conditions. Cells were then washed three times with PBS and resuspended with 50 µl of 80% glycerol. Approximately, 10 µl of this cell suspension was spotted on the microscopic glass slide and covered with a coverslip.

### Rhodamine-phalloidin staining for F-actin

Staining of polymerized actin filaments (F-actin) in *C. albicans* mutant strains was performed with rhodamine-phalloidin (300 U) at a dilution of 1:30 following standard protocols and used for microscopy (28).

### Transcript levels analysis using qPCR

Isolation of RNA and cDNA synthesis were done from mid-log phase cells as described previously (29). The qPCR was performed using SYBR green PCR master mix (Kapa biosystems) and gene specific RT primers. The expression of genes at mRNA levels were analysed using the comparative *C_t_*method. *GAPDH* was used as the internal control.

### Pull-down assays and western blots

To study whether CaEri1 interacts with CaGpi2, *CaERI1* was tagged with V5 at the C-terminus in BWP17-*CaGPI2-6X-His* (to express CaGpi2 tagged with His_6_ at its C-terminus). Secondary culture of BWP17-*CaGPI2-6X-His/CaERI1-V5* strain was cultured till O.D_600 nm_ reached between 1 to 2. Cells were lysed in lysis buffer (10 mM HEPES-Na pH 7.5, 200 mM (NH_4_)_2_SO_4_, 5 mM MgSO_4_, 5 mM EDTA, 0.1% NP-40 substitute, 200 mM NaCl, 0.5% Triton-X, 10% glycerol) with glass beads, vortexed for on ice (1 min x 10 rounds). The supernatant obtained after centrifugation at 3000 rpm (4 °C) was incubated overnight with Ni^+2^-NTA beads at 4 °C. The unbound supernatant was removed and beads washed with lysis buffer (1 ml x 6 rounds). Bound proteins were eluted in SDS-PAGE sample buffer at 95 °C and resolved on 12% SDS-PAGE. Proteins were transferred onto a PVDF membrane and probed with one of the following antibodies (4 °C, overnight): Anti-His (#SC-8036; Santa Cruz biotechnology) at 1:700 dilution, Anti-V5 (#13202, Cell Signaling Technology) at 1:1000 dilution, anti-Ras (Clone RAS10, #05-516, Merck-Millipore) at 1:1000 dilution and anti-G6PDH (A9521, Merck-Millipore) at 1:8000 dilution. HRP-conjugated secondary antibody (#114068001A, Genei) at a dilution of 1:5000 and 1: 8000 was added and the membranes were incubated at room temperature for 2 h. The blots were probed using ECL reagent (Luminata Forte, WBLUF0100, Merck-Millipore).

### Acceptor-photobleaching Förster resonance energy transfer (AP-FRET)

CaEri1-V5 was expressed in BWP17 as well as in *CaGPI2*-*6X-HIS* strains. CaRas1 was detected by primary anti-Ras antibodies while 6X-His and V5 tagged proteins were detected by primary anti-6X-His and anti-V5 antibodies, respectively. Appropriate secondary antibodies were used (Goat anti-mouse IgG-FITC and Goat anti-rabbit IgG-TRITC) to generate the fluorescent FITC-TRITC donor-acceptor pairs. *CaGPI2*-*6X-HIS*/*CaRAS1* and *CaERG11*-*6X-HIS*/*CaRAS1* strains were used as the positive and negative control strains, respectively (8). AP-FRET was performed using a Nikon A1R HD25 confocal microscope using a previously standardized protocol (8). Briefly, FITC was excited with a 488 nm HeNe laser and emission was collected using a 500-530-nm band-pass filter. TRITC was excited with a 543 nm HeNe laser and emission was collected using a 550-590 nm long-pass filter. A neighboring cell was monitored as a control before and after photobleaching to track for non-specific effects. The TRITC label was photobleached by repeated irradiation with a 543 nm HeNe laser. Images of fluorescence emission from the cell in the FITC and TRITC channels were again captured (pre- and post-bleach). No bleed-through was observed in the TRITC-channel for cells labelled only with FITC-conjugated antibodies alone. No non-specific excitation of TRITC by the 488 nm laser was observed in cells labelled with TRITC-conjugated antibodies alone. FRET efficiency was calculated as follows:

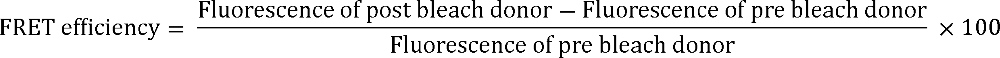

Background subtraction was done by selecting an ROI outside the cell. FRET observations were obtained for 8 cells in each set and quantified using Nikon NIS element 4000 AR analysis software.

### Determination of fungal burden in *G. mellonella* larvae

The sixth instar larval stage of *G. mellonella*, having length of ∼2 cm and ∼0.3-0.4 gm, without any melanisation or grey markings were used for experiments (30). To assess the fungal burden in the larvae, 5 larvae of in each group were taken for the experiments. *C. albicans* cells from secondary cultures were harvested by centrifugation (5000 rpm, room temperature, 5 min) and washed with sterile PBS, pH 7.2. Cell density was determined by a hemocytometer. Cells at a density of 1× 10^6^ cells/20 µl were inoculated through the last left pro-leg of the larvae using 0.5 ml sterile insulin syringe (HMD, India) for the infected groups. Larvae were incubated at 30 °C for 24 h. Three random larvae from each group were homogenized individually by adding 1 ml of sterile PBS, pH 7.2. Appropriate dilutions were made and plated on YEPD plates containing kanamycin (100 µg/ml) to prevent bacterial growth. The plates were incubated at 30 °C for 24 h. The fungal burden was determined as colony forming units (CFU)/ml as given below:

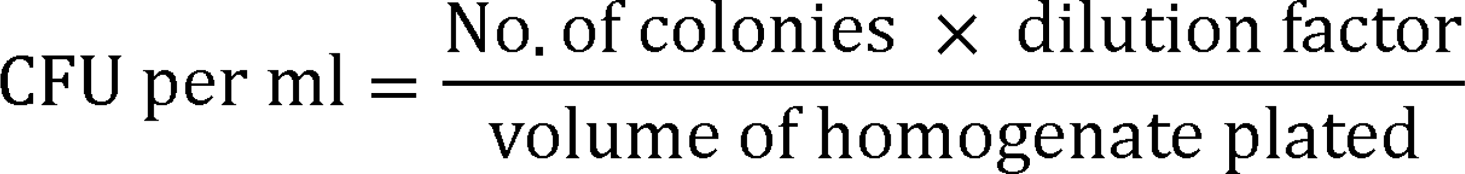

### *G. mellonella* larval survival assay

For these assays, four groups of larvae were used, and each infection group consisted of 10 larvae. Two groups were the control groups: 1) where larvae were not touched (untouched) and 2) the larvae were injected with PBS only. The two other two groups were infected with either BWP17-*URA3*, *CaERI1Hz-URA3* or *CaERI1* reintegrant strains at 5× 10^5^ cells/20 µl as described above. Larvae were incubated at 37 °C after inoculation and larval death was monitored every 6 h.

### Statistical analysis

Paired Student’s t-test was performed to analyse the statistical significance. Symbols, *, ** and *** denote *P*-values < 0.05, < 0.01 and < 0.001, respectively. GraphPad Prism 5.0 software was used for the analysis.

## Acknowledgements

Parts of this work were funded by grants to SSK from Department of Biotechnology (DBT) India (BT/PR29186/BRB/0/1726/2018) and Science and Engineering Research Board, India (SERB:CRG/2020/001649). Facilities and research at the School of Life Sciences are supported by grants from UGC-CAS, UGC-RNW, DBT-BUILDER and DST-FIST. SCS and MB were supported by Senior and Junior Research fellowships from University Grants Commission, India. SCS also received salary from the SERB grant of SSK. YK and UY received JRF and PDF fellowships, respectively, from Indian Council of Medical Research. We thank Prof. David E. Levin for 1788 and DL2524 strains, Dr. Cheryl Gale for *URA3-pMET3-GFP* vector, Prof. Alistair JP Brown for *pACT1-GFP* vector and Prof. Aaron Mitchell for the BWP17 strain. *G. mellonella* insect larvae were a kind gift from Prof. Rakesh. K. Seth, University of Delhi, India. Confocal images were recorded at the Central Instrumentation Facility, SLS, JNU. The funders had no role in study design, data collection and analysis, decision to publish, or preparation of the manuscript.

## Author Contributions

**Sneha Sudha Komath:** Conceptualization, Funding acquisition, Supervision, Visualization, Writing-Original Draft, Resources, Formal analysis; **Subhash Chandra Sethi:** Methodology, Investigation, Validation, Formal analysis, Writing-Original Draft; **Monika Bharati:** Methodology, Investigation, Validation, Formal analysis, Writing-Original Draft; **Usha Yadav:** Methodology, Investigation; **Yatin Kumar:** Methodology, Investigation, Formal analysis. All authors read and finalized the final draft.

