## Supplementary Figures and Table I for "The ER-resident Ras Inhibitor 1 (Eri1) of *Candida albicans* inhibits hyphal morphogenesis via the Ras-independent cAMP-PKA pathway": Sethi et al. supplementary files.docx

^2^Present address: Genetics Branch, Center for Cancer Research, National Cancer Institute, National Institutes of Health, Bethesda, MD, USA

**Running Title:** Role of *C. albicans* Eri1

**Keywords:** *Candida albicans* Eri1, GPI-*N*-acetylglucosaminyltransferase*,* hyperfilamentation, cAMP-PKA signaling, inter-subunit regulation, *Galleria mellonella* infection model

**Supplementary Information:**

**Figure S1**

**Figure S2**

**Figure S3**

**Figure S4**

**Table S1**

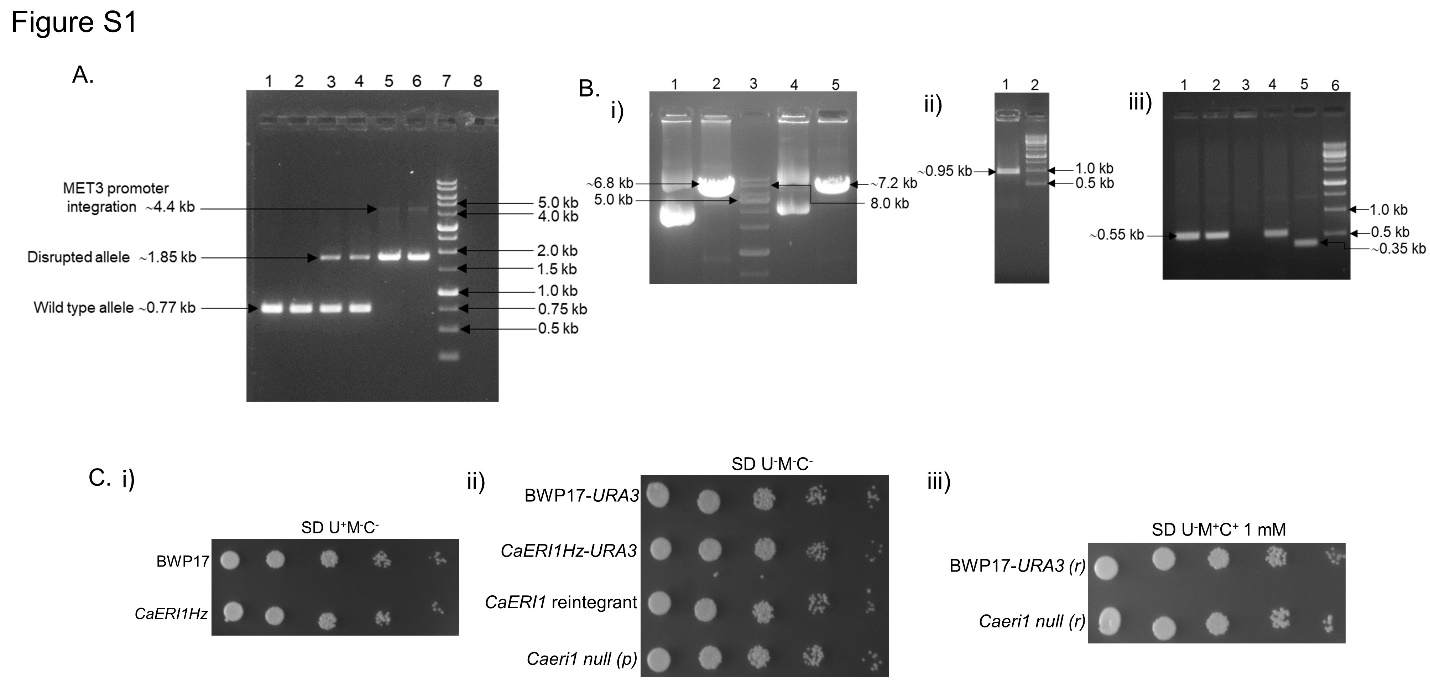

**Figure S1:** **Generation, confirmation and growth of *Caeri1* mutant strains in SD agar medium. A. Confirmation of *CaERI1Hz* and *Caeri1* conditional null mutant.** The *HIS1* selection marker was used to generate the heterozygous mutant while the conditional null mutant was generated by replacing the native promoter of the existing *CaERI1* allele with a regulatable *MET3* promoter in the *CaERI1Hz* background. The mutants were confirmed by *CaERI1* locus specific flanking forward and reverse primers using the genomic DNA as template. Lane 1,2: BWP17; Lane 3,4: positive colonies of the *CaERI1Hz* mutant; Lane 5,6: positive colonies of *Caeri1* *null* mutant; Lane 7: 1 kb DNA ladder; Lane 8: negative control. **B. Confirmation of the *CaERI1Hz-URA3* and *CaERI1* revertant strain**. (i) Digestion of p*ACT1-CaERI1* and p*ACT1-GFP* with StuI. Lane 1: undigested p*ACT1-CaERI1*, Lane 2: StuI digested p*ACT1-CaERI1*; Lane 3: 1 kb DNA ladder; Lane 4: undigested p*ACT1-GFP*, Lane 5: StuI digested p*ACT1-GFP*. (ii) The StuI digested p*ACT1-GFP* plasmid was used to transform the *CaERI1Hz* strain. Confirmation of the integration of p*ACT1-GFP* at *RPS1* locus to generate *CaERI1Hz-*p*ACT1-GFP* (*CaERI1Hz-URA3*) strain using PCR with *GFP* specific FP and *RPS1* specific RP. Lane 1: positive colony; and Lane 2: 1kb DNA ladder. (iii) The StuI digested p*ACT1-CaERI1* plasmid was used to transform the *CaERI1Hz* strain to generate *CaERI1Hz*/p*ACT1*-*CaERI1* (*CaERI1* revertant) strain. *CaERI1* specific FP and *RPS1* specific RP were used to confirm the *CaERI1* revertant strain by PCR to amplify the band of interest from genomic DNA of the transformants, whereas *CaERI1* specific FP and RP were used to amplify the positive control from BWP17 genomic DNA. Lane 1-2 and 4: Positive colonies; Lane 3: negative colony; Lane 5: positive control; Lane 6: 1 kb DNA ladder. **C.** ***Caeri1* mutants do not exhibit growth defects in solid medium** (i-ii) None of the strains showed significant growth defects on SD-agar plates. *CaERI1Hz* and BWP17 strains were grown on media containing uridine. BWP17-*URA3*, *CaERI1Hz -URA3, Caeri1 null (p),* and *CaERI1 reintegrant* were grown on SD media in the absence of Met/Cys (Met^-^Cys^-^). BWP17-*URA3* and *Caeri1 null (r)* were also grown on SD media containing 1 mM Met/Cys (Met^+^Cys^+^).

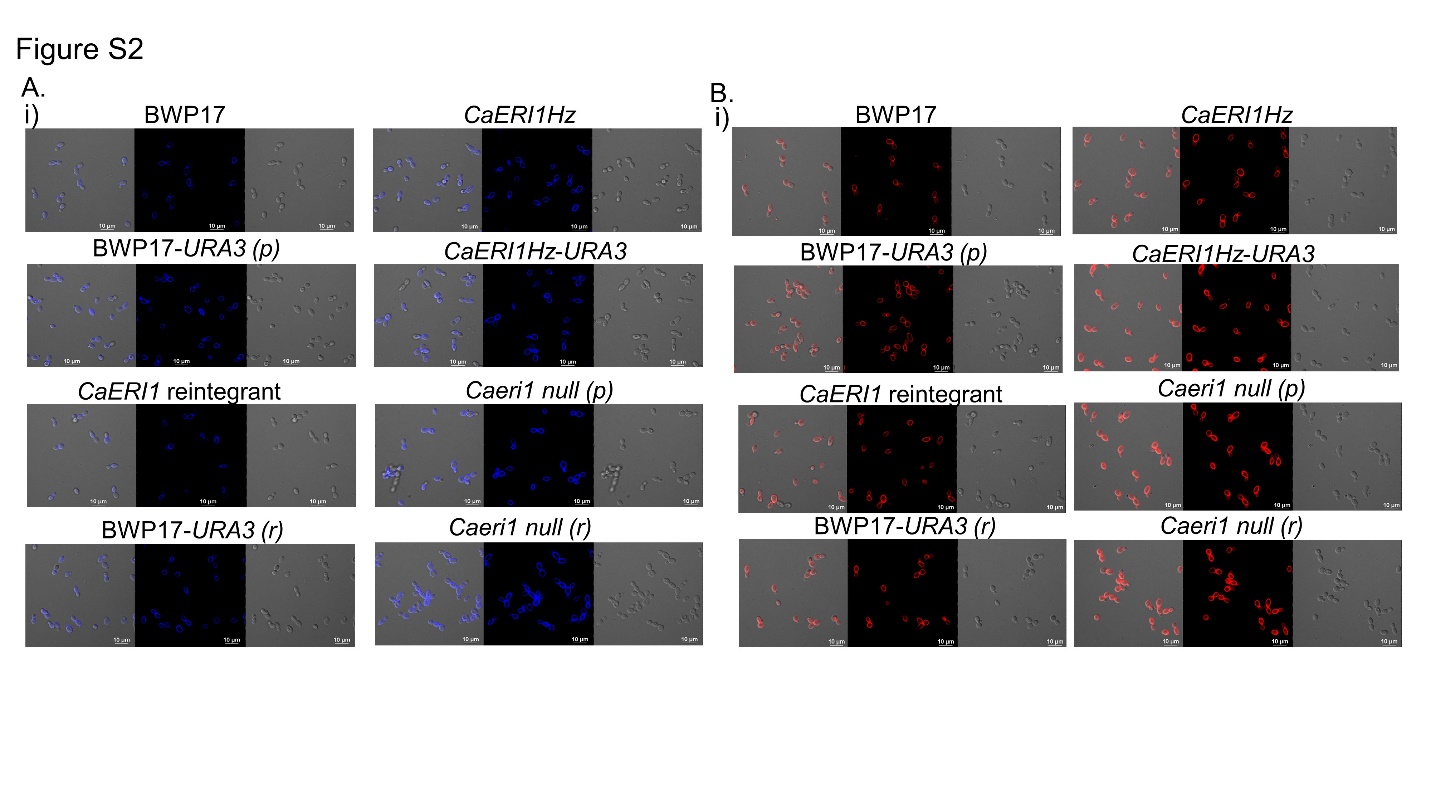

**Figure S2: Staining of cell wall components, chitin and β-glucan using Calcofluor White (CFW) and Congo Red (CR), respectively. A Staining with CFW for chitin.** (i-iii) The indicated *C. albicans* strains were stained with 100 µg/ml CFW for 30 min and images were captured using a Nikon Eclipse TiE fluorescence microscope on DAPI channel. **B.** **Staining with CR for β-glucan.** (i-iii) The mentioned strains were stained with 100 µg/ml CR for 30 min and images were captured using a Nikon Eclipse TiE fluorescence microscope on TRITC channel. These experiments were repeated three times with independent cultures.

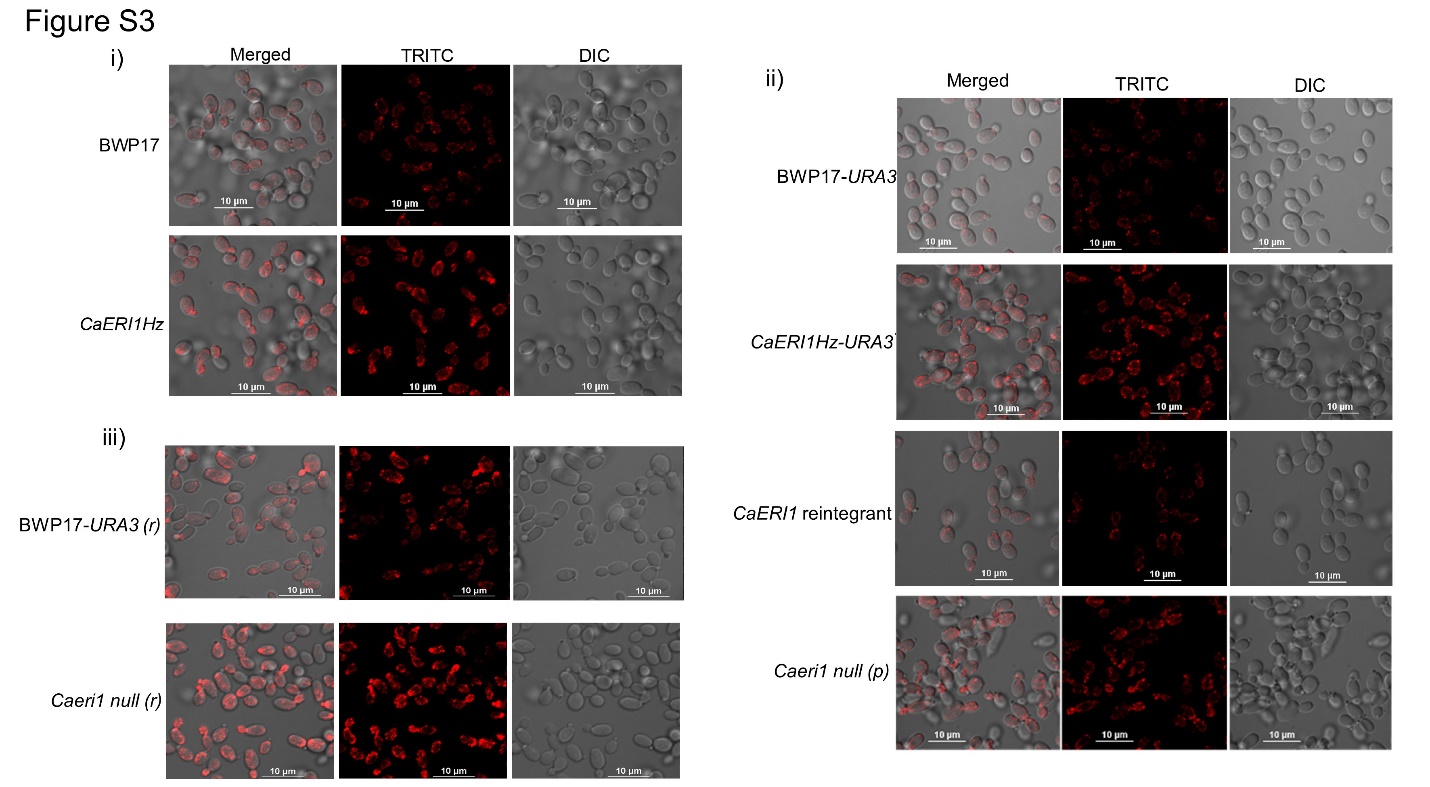

**Figure S3: F-actin staining using rhodamine-phalloidin in *Caeri1* mutant strains.** (i-iii) The *Caeri1* mutant strains were stained with rhodamine-phalloidin as per the protocol described in the ‘Experimental Procedures’ section. The stained cells were visualized on TRITC channel under a Nikon A1R confocal microscope. These experiments were done three times with independent cultures for confirmation.

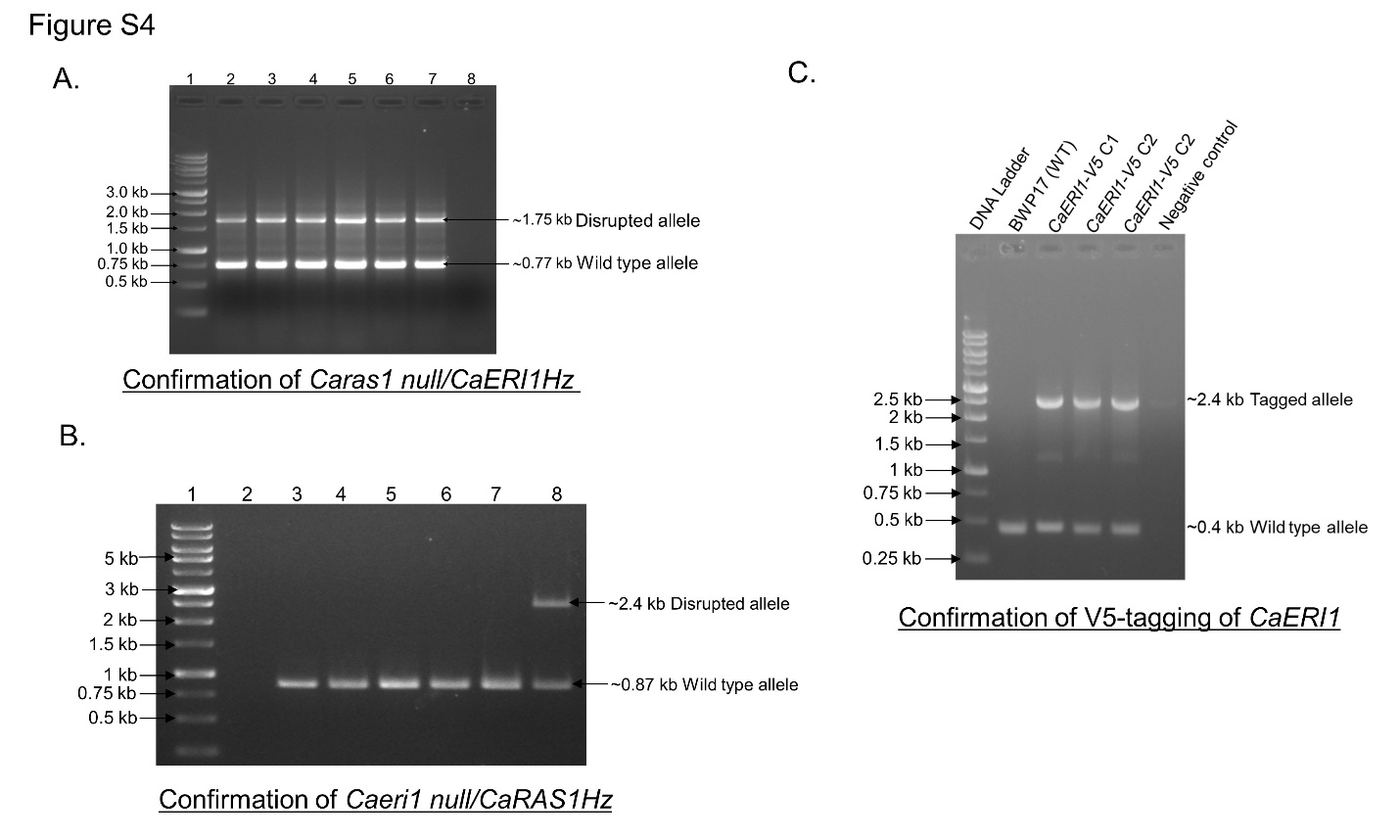

**Figure S4: Confirmation of *Caeri1* mutant strains. A. Confirmation of *Caras1 null/CaERI1Hz* strain.** One allele of *CaERI1* was deleted in *Caras1 null* strain by using *URA3* selection marker through PCR mediated disruption approach. The *URA3* cassette was amplified using primers such as CaERI1 URA3 FP and CaERI1 URA3 RP (Table S1). The amplicon was used to transform the *Caras1 null* strain. The transformants were screened by setting PCR with *CaERI1* locus specific flanking forward and reverse primers. Lane 1: 1 kb DNA ladder, Lane 2-7: positive colonies (Band size of ~0.77 kb from the wild type allele; band size of ~1.75 kb from the disrupted allele), Lane 8: negative control. **B.** **Confirmation of *Caeri1 null/CaRAS1Hz* strain.** One allele of *CaRAS1* was disrupted in *Caeri1null* strain by using *URA3* selection marker. The *URA3* cassette was amplified using primers such as CaRAS1 ARG4 FP and CaRAS1 ARG4 RP**.** The amplicon was then used to transform the *Caeri1 null* strain and the obtained transformants were screened by performing PCR with *CaRAS1* locus specific flanking forward and reverse primers. Lane 1: 1 kb DNA ladder, Lane 2: negative control, Lane 3-7: negative colonies, Lane 8: a positive colony (Band size of ~0.87 kb from the wild type allele; band size of ~2.4 kb from the disrupted allele). **C.** **Confirmation of *CaERI1-V5* tagging.** The V5 epitope tagging of CaEri1 at its C-terminus was done using the *ARG4* selection marker. The *ARG4* marker was amplified using CaERI1-V5 ARG4 FP and CaERI1 ARG4 RP. The amplicon was then transformed to BWP17, colonies obtained were screened by PCR with CaERI1 internal FP and CaERI1 locus specific RP. Lane 1: DNA ladder, Lane2: BWP17, Lane 3-5: positive colonies (Band size of ~ 0.4 kb wild type allele and band size of ~ 2.4 kb tagged allele), Lane 6: Negative control.

**Table S1: List of primers used in the study**

| **Primer name** | **Primer sequence** |
| --- | --- |
| FPCaGPI2-ARG4 | 5'-ATGGAAGAAATACATATAAGCTCTTCCATTTTGCAAGATA GTGATTCAGCTCCCCCAAACCATTTTCAGTGTGGAATTGTGA GCGGATA-3' |
| RPCaGPI2-ARG4 | 5'-TCAGCTTTGTATACTTGACTTCATTAACTTTGGTTTCGCC ACATCCCATGGTCCTTGTATCTCGTTTTTTTTCCCAGTCAC GACGTT-3' |
| FPCaGPI2-UP | 5'-AAGTATTCTATATCTTGTGTTTTGTGGGTCGTGTTT-3' |
| RPCaGPI2-DOWN | 5'-CACTGAAAAGAAAAGTGTATATTATCTTCTATAAGG-3' |
| FPCaGPI2-HindIII | 5'-GCGAAGCTTATGGAAGAAATACATATAAGCTCT-3' |
| RPCaGPI2-NheI | 5'-GCGGCTAGCTCAGCTTTGTATACTTGACTTCAT-3' |
| FPCaGPI15-HIS1 | 5'-ATGTCCTCATCGAAAAATTATAAATTGGAAATTTCCCCT AGTAGTATCAACCAAACTAGTGCCACAGAATCGCGGGGA TCCTGGA GGATGAG-3' |
| RPCaGPI15-HIS1 | 5'-CTAAATAACTTGTTTTAAACCTTGACCAGGCACTCTTCTCCA ATAGCGTTTTGTGCTTCCAAATAGTAACGGAATATTTATGAG AAACT-3' |
| Flanking FPCaGPI15 | 5'-TGGTAACCATTCATCAATCATATAG-3' |
| Flanking RPCaGPI15 | 5'-GGTTTATACATAGATTTTATATACATTG-3' |
| FPCaGPI19-HIS1 | 5'-ATGATATTCCATTTTAACCAAAAAGAGAAAGCAGATTCAAG TAAATGTAAAACAATACATGTAAGAAAGACGGGGATCCTGG AGGATGAG-3' |
| RPCaGPI19-HIS1 | 5'-TCATTCATATAGAACATCATTCATAATGTAATAGGCAAA TCC CAAAC ACCACTTGGTGCCTTAGAATACGGAATATTTA TGAGA AACT-3' |
| FPCaGPI19-ARG4 | 5'-ATGATATTCCATTTTAACCAAAAAGAGAAAGCAGATTCAAG TAAATGTAAAACAATACATGTGGAATTGTGAGCGGAAG-3' |
| RPCaGPI19-ARG4 | 5'-TCATTCATATAGAACATCATTCACTAATGTAATAGGCAAA TCCCAAA CACCACTTGGTGCCTTTCCCAGTCACGACGTT-3' |
| FPCaGPI19-HindIII | 5'-GCGAAGCTTATGATATTCCATTTTAACCAAAAAG-3' |
| RPCaGPI19-NheI | 5'-GCGGCTAGCTCATTCATATAGAACATCATTCACT-3' |
| FPCaERI1-HIS1 | 5'-ATGCGTTCACAATCATTGCCATTACCATTAATAGAATCACC AAACTCAATAGATTTAGACCCCATCAATCCGGGGATCCTGGAGGATGAG-3' |
| RPCaERI1-HIS1 | 5'-TTACCCTTGAATACCTTTAGAATGACGGAATAATTTTAAC CCACTCCATGATACAATGGCCCACCACCAACGGAATATTTA TGAGAAACT-3' |
| Flanking FPCaERI1 | 5'-AGTGAAAGAAGGACGTGGTTTAC-3' |
| Flanking RPCaERI1 | 5'-GGGTATTGAGCTAAAAGAGGAGGT-3' |
| FPCaERI1-BamHI | 5'-GCGGGAATCCATGCGTTCACAATCATTTAGC-3' |
| RPCaERI1-MluI | 5'-GCGACGCGTTTACCCTTGAATACCTTTAGA-3' |
| RPCaRPS1 | 5'-AATAGAGAGAAACTATATTATACAC-3' |
| FPCaGPI2-HIS1 | 5'-ATGGAAGAAATACATATAAGCTCTTCCATTTTGCAA GATAGTGATTCAGCTCCCCCAAACGGGGATCCTGGAGGATGAG-3' |
| RPCaGPI2-HIS1 | 5'-TCAGCTTTGTATACTTGACTTCATTAACTTTGGTTTCGCCA CATCCCATGGTCCTTGTACGGAATATTTATGAGAAACT-3' |
| FP CaERI1-URA3 | 5'-GCTCTATGTCACAATACATTAACAAAGTCAATCACTACAT GTTTCCCTGACTAGTCTAGAAGGACCACCTTTGATTG-3' |
| RPCaERI1-MET3 | 5'-CTAAATCTATTGAGTTTGGTGATTCTATTAATGGTAATGGC AATGATTGTGAACGCATCATTTTAATAAACGCGGATCC-3' |
| FPCaGPI15-ARG4 | 5'-ATGTCCTCATCGAAAAATTATAAATTGGAAATTTCCCCTAG TATCAACCAAACTAGTGCCACAGAATCGTGTGGAATTGTGAGGGGAAG-3' |
| RPCaGPI15-ARG4 | 5’-TCTAAACATTACTTGTTTATTTCTATTGGGTTTTTTTATTAT AAAGGTTTTGTTTTTAACTTAAAATTGATTTTAAATT-3’ |
| FPCaERI1-URA3 | 5’-ATGCGTTCACAATCATTGCCATTACCATTAATAGAATCACCAAACTCAATAGATTTATAATAGGAATTGATTTGGATGG-3' |
| RPCaERI1-URA3 | 5’-TTACCCTTGAATACCTTTAGAATGACGGAATAATTTTAACCCACTCCATGATACAATTATAATTGGCCAGTCTTTTTC-3' |
| FPCaERI1-ARG4 | 5’-ATGAGAATTTGGCGGTATACGGATATTTGCTACTAATATCAACATGGATGTTATTCATTTGTGGAATTGTGAGCGGATA-3’ |
| RPCaERI1-ARG4 | 5’-ATACTACATAGATACACCATAACTTTATTATATAGGAATCAATAGTTTCGAAGAAAATTGTTTCCCAGTCACGACGTT-3’ |
| FPCaRAS1-URA3 | 5’ATGTTGAGAGAATATAAATTAGTTGTTGTTGGAGGTGGTGGTGTTGGTAAATCCGCTTAATAGGAATTGATTTGGATGG-3’ |
| RPCaRAS1-URA3 | 5’- TCAAACAATAACACAACATCCATTCTTTGATTTAGAGCTAGATTGTTTTTGACCAGAATATAATTGGCCAGTCTTTTTC -3’ |
| FlankingFPCaERI1-V5 | 5’- GCGGTATACGGATATTTGCTACTAATATCA-3’ |
| FlankingRPCaERI1-V5 | 5’- GAATCTTAGATTATGCCAGTTGATCCTTTG-3’ |
| RT FPCaGAPDH | 5'-CAGCTATCAAGAAAGCTTCTG-3' |
| RT RPCaGAPDH | 5'-GATGAGTAGCTTGAACCCAA-3' |
| RT FPCaGPI2 | 5'-GGCCAATAGCATTTCTAAC-3' |
| RT RPCaGPI2 | 5'-CCATCACAAAAACAGACAAA-3' |
| RT FPCaGPI15 | 5'-CCACGATTATGGGCAGGTTA-3' |
| RT RPCaGPI15 | 5'-CGCGGTAAGAATTCTGGAAA-3' |
| RT FPCaGPI19 | 5'-CAAGAAGAAGAAGAAGGAGAA-3' |
| RT RPCaGPI19 | 5'-AAACACCACTTGGTGCCTTA-3' |
| RT FPCaERI1 | 5'-ACAGTCATCGCCAACATCCT-3' |
| RT RPCaERI1 | 5'-ATGGCCCACCACCAACATAC-3' |
| RT FPCaEFG1 | 5'-AGCTCCAATACAAGTTGCA-3' |
| RT RPCaEFG1 | 5'-TCCCAATTTACCAGCAGACAAGA-3' |
